# Decoding the transcriptomic signatures of psychological trauma in human cortex and amygdala

**DOI:** 10.1101/2024.10.23.619681

**Authors:** Emily M. Hicks, Carina Seah, Michael Deans, Seoyeon Lee, Keira J.A. Johnston, Alanna Cote, Julia Ciarcia, Akash Chakka, Lily Collier, Paul E. Holtzheimer, Keith A. Young, Traumatic Stress Brain Research Group, John H. Krystal, Kristen J. Brennand, Eric J. Nestler, Matthew J. Girgenti, Laura M. Huckins

## Abstract

Psychological trauma has profound effects on brain function and precipitates psychiatric disorders in vulnerable individuals, however, the molecular mechanisms linking trauma with psychiatric risk remain incompletely understood. Using RNA-seq data postmortem brain tissue of a cohort of 304 donors (N=136 with trauma exposure), we investigated transcriptional signatures of trauma exposures in two cortical regions (dorsolateral prefrontal cortex, and dorsal anterior cingulate cortex) and two amygdala regions (medial amygdala and basolateral amygdala) associated with stress processing and regulation. We focused on dissecting heterogeneity of traumatic experiences in these transcriptional signatures by investigating exposure to several trauma types (childhood, adulthood, complex, single acute, combat, and interpersonal traumas) and interactions with sex. Overall, amygdala regions were more vulnerable to childhood traumas, whereas cortical regions were more vulnerable to adulthood trauma (regardless of childhood experience). Using cell-type-specific expression imputation, we identified a strong transcriptional response of medial amygdala excitatory neurons to childhood trauma, which coincided with dysregulation observed in a human induced pluripotent stem cell (hiPSC)-derived glutamatergic neurons exposed to hydrocortisone. We resolved multiscale coexpression networks for each brain region and identified modules enriched in trauma signatures and whose connectivity was altered with trauma. Trauma-associated coexpression modules provide insight into coordinated functional dysregulation with different traumas and point to potential gene targets for further dissection. Together, these data provide a characterization of the long-lasting human encoding of traumatic experiences in corticolimbic regions of human brain.

## INTRODUCTION

There is a powerful link between trauma and mental health; traumatic stress is a major precipitating factor in the development of several psychiatric disorders, particularly mood and anxiety disorders, such as post-traumatic stress disorder (PTSD) and major depressive disorder (MDD). Yet, the molecular mechanisms linking traumatic stress to psychiatric disorder risk are only beginning to be elucidated^1–3^. Traumatic experiences are processed through corticolimbic circuits^4^ and mount a whole body-endocrine response through glucocorticoid (GC) signaling^5,6^. Both circuit and endocrine systems employ robust negative feedback mechanisms to return the organism to homeostasis, however, dysfunction in these mechanisms, for example, undermodulation of limbic circuits by prefrontal cortex^7^ and GC hyper-/hypo-sensitivity^8^ precipitate prolonged psychiatric symptoms such as intrusions, avoidance, altered mood or arousal as a consequence.

It is critical to examine the contexts in which traumas occur. Since corticolimbic stress circuitry is molded in childhood and adolescence through critical periods of rapid synaptic growth, pruning and myelination^9–11^, trauma during development has lasting effects of emotional and cognitive processing^12–14^, and leads to a greater risk for psychiatric disorder in adulthood^12^. Different trauma types are also associated with differential risk for PTSD^15^. We and others have shown that different types of traumas predispose to disorder at different rates; for example, sexual traumas and childhood traumas are associated with increased rates of post-traumatic stress disorder (PTSD) compared to other traumas, even in the context of exposure to combat trauma or disasters^16–18^. Moreover, exposure to traumatic stress likely has different outcomes according to sex and gender, through a myriad of mechanisms both psychosocial (e.g., access to social support, stigmatization, and exposure to different types of traumas) and physiological (e.g., through interaction between stress hormones and reproductive hormones such as estradiol^19,20^).

Recent transcriptional studies of human postmortem brain have demonstrated transcriptomic dysregulation characteristics of traumatic stress-related psychiatric disorders in both a brain region- and cell type-specific manner, highlighting the importance of corticolimbic regions and several cell types^21–27^. Where human postmortem brain samples capture the breadth of lived experiences, cultured neurons derived from human-induced pluripotent stem cells (hiPSCs) isolate cell type specific effects of exposure in a naïve state. With this model, we have previously characterized a robust transcriptional response of hiPSC-derived glutamatergic neurons to hydrocortisone (HCort) ^28^. Evidence suggests that trauma-exposed patients with psychiatric symptoms may represent a distinct subtype of disorder (aside from PTSD, for which trauma exposure is a requirement) ^13,29^. We hypothesize that trauma per se produces a lasting transcriptional “scar” that is agnostic of diagnosis. To date, no published studies have examined the transcriptomic impacts of psychosocial trauma, and dissected how the encoding of various types of traumas accumulate in the human brain.

In this study, we investigate the lasting transcriptional encoding of trauma in the human dorsolateral prefrontal cortex (DLPFC), dorsal anterior cingulate cortex (dACC), and medial amygdala (MeA) and basolateral amygdala (BLA) in a large cohort of postmortem human brain samples of individuals with extensive trauma history phenotyping. We first characterize the transcriptional signatures and coexpression relationships of exposure to and accumulation of trauma throughout the lifespan in a brain region- and cell type-specific manner. Next, we explore several dimensions of trauma, including developmental period, trauma type, and trauma load, and highlight convergent and divergent mechanisms in their transcriptional signatures. Using hiPSC-derived excitatory neurons, we explore convergent transcriptional dysregulation across childhood trauma and hydrocortisone. Finally, we identify sex-specific gene expression interactions and investigate potential mechanisms underlying sex differences (e.g., trauma type, neuronal sex hormone interactions).

## METHODS

### Human postmortem cohort

Human autopsy brain tissue samples from 304 individuals were donated at time of autopsy through U.S. medical examiners’ office and from the VA National PTSD Brain Bank and Traumatic Stress Brain Research group. Psychiatric histories and demographic information were obtained by postmortem psychological autopsies conducted by clinicians with informants best acquainted with the individual and diagnostic review of medical records. Sex was defined as sex assigned at birth: 115 total donors were female, 189 were male. Race was assigned by clinician during psychological autopsy: donors represented a mix of White, Black and Hispanic. Mean age of death for all donors was 46.75 years. A full description of sample processing and sequencing can be found elsewhere^23^.

### Trauma phenotyping

We catalogued each recorded lifetime traumatic exposure by literature-derived trauma categories (combat vs interpersonal traumas) and age group at which the traumatic event occurred: childhood (<20 y) or adulthood (>20y). Trauma histories were coded in three independent replicates (by 3 independent researchers) to reach a consensus coding for each donor. Each donor was assigned one value or category for each trauma variable (**Table 1**) and summarized across the cohort demographics (**Table 2**).

**Table 1:** Trauma variables and category descriptions for phenotyping donors from trauma history information.

| **Trauma Variable** | **Trauma Variable Category** | **Description** |
| --- | --- | --- |
| TraumaAny | Any trauma | At least one trauma exposure |
|  | No trauma | 0 reported lifetime traumas |
| Trauma Count | NA | Cumulative number of trauma exposures |
| Childhood Adulthood | Childhood only | At least one reported trauma in childhood (<20 years old) and 0 reported adulthood ( 20+ years old) traumas |
|  | Adulthood only | 0 reported traumas in childhood (<20 years old) and at least one reported adulthood ( 20+ years old) trauma |
|  | Both | At least one reported trauma in childhood (<20 years old) and at least reported trauma in adulthood ( 20+ years old) |
|  | No trauma | 0 reported lifetime traumas |
| Childhood Count | NA | Cumulative number of trauma exposures in childhood |
| Adulthood Count | NA | Cumulative number of trauma exposures in adulthood |
| Single Complex | Single only | At least one reported acute trauma (e.g. accident, assault) and 0 reported complex traumas (e.g. physical abuse, sexual abuse, neglect) |
|  | Complex only | 0 reported acute traumas (e.g. accident, assault) and at least one reported complex traumas (e.g. physical abuse, sexual abuse, neglect) |
|  | Both | At least one reported acute trauma (e.g. accident, assault) and at least one reported complex trauma (e.g. physical abuse, sexual abuse, neglect) |
|  | No trauma | 0 reported lifetime traumas |
|  | (Exclusions) | At least one reported combat trauma |
| Combat Interpersonal | Combat only | At least one reported combat exposure and 0 reported interpersonal traumas (e.g., abuse, assault) |
|  | Interpersonal only | 0 reported combat exposures and at least one reported interpersonal trauma (e.g., abuse, assault) |
|  | Both | At least one reported combat exposure and at least 1 reported interpersonal traumas (e.g., abuse, assault) |
|  | No trauma | 0 reported lifetime traumas |
| Top Bottom Quartile | Top | Top quartile of cumulative trauma load |
|  | Bottom | Bottom quartile of cumulative trauma load (trauma load > 0 ) |
|  | No trauma | 0 reported lifetime traumas |

**Table 2:** Sample size (N), sex and race and average age for samples in each trauma variable category. AFR=African American, EUR=European American, HISP=Hispanic, NT=neurotypical controls, MDD=major depressive disorder, PTSD= posttraumatic stress disorder

| **Trauma Measure** | **Trauma Category** | **N** | **F** | **M** | **AA** | **EUR** | **HISP** | **NT** | **MDD** | **PTSD** | **Avg, Age** |
| --- | --- | --- | --- | --- | --- | --- | --- | --- | --- | --- | --- |
| Trauma Any | Any Trauma | 137 | 70 | 67 | 15 | 121 | 1 | 2 | 31 | 104 | 42.1 |
|  | No Trauma | 167 | 45 | 122 | 35 | 129 | 3 | 92 | 75 | 0 | 48.2 |
| Childhood Adulthood | Childhood Only | 47 | 24 | 23 | 7 | 39 | 1 | 1 | 13 | 33 | 43.8 |
|  | Adulthood Only | 45 | 18 | 27 | 5 | 40 | 0 | 1 | 6 | 38 | 42.2 |
|  | Both | 22 | 14 | 8 | 2 | 20 | 0 | 0 | 2 | 20 | 39.3 |
|  | No Trauma | 162 | 45 | 117 | 35 | 124 | 3 | 89 | 73 | 0 | 47.8 |
| Single Complex | Single Only | 17 | 8 | 9 | 0 | 17 | 0 | 0 | 4 | 13 | 41.0 |
|  | Complex Only | 55 | 30 | 25 | 7 | 47 | 1 | 1 | 19 | 35 | 46.1 |
|  | Both | 45 | 31 | 14 | 3 | 42 | 0 | 0 | 8 | 37 | 38.5 |
|  | No Trauma | 167 | 45 | 122 | 35 | 129 | 3 | 92 | 75 | 0 | 48.2 |
| Combat Interpersonal | Combat Only | 19 | 0 | 19 | 5 | 14 | 0 | 1 | 0 | 18 | 39.5 |
|  | Interpersonal Only | 99 | 65 | 34 | 10 | 88 | 1 | 1 | 28 | 70 | 42.9 |
|  | No Trauma | 162 | 45 | 117 | 35 | 124 | 3 | 89 | 73 | 0 | 47.8 |
| Top Bottom Quartile | Top | 34 | 24 | 10 | 1 | 32 | 1 | 0 | 4 | 30 | 41.7 |
|  | Bottom | 35 | 11 | 24 | 7 | 28 | 0 | 2 | 7 | 26 | 41.2 |
|  | No Trauma | 167 | 45 | 122 | 35 | 129 | 3 | 92 | 75 | 0 | 48.2 |

### Multiple correspondence analysis of trauma phenotypes

To determine the level of similarity in the trauma variable categories within our cohort, we ran multiple correspondence analysis (MCA) on the donor trauma profiles using factoMineR package ^30^. This analysis can be thought of as a generalization of principal component analysis for categorical data and finds the dimensions associated with the largest sources of variation in the data. We plotted each trauma variable category and its association with MCA dimensions to demonstrate the relatedness of our definitions within our cohort.

### RNA-seq preprocessing and quality control

RNA transcript counts were log2 transformed and normalized for library size. We removed lowly expressed genes by requiring at least 20 counts per gene in at least 2 samples and transcripts from sex chromosomes and mitochondrial RNA then voom normalized for linear modeling^31^.

### Gene differential expression and trauma-leading gene analysis

We used surrogate variable analysis (SVA)^32^ to calculate surrogate variables (SVs) that capture measured and latent sources of variation in gene expression for each brain region, while preserving the effects of Trauma_Count_, sex and diagnosis. We calculated 23, 22, 25 and 28 SVs, respectively, for the DLPFC, dACC, BLA and MeA, and verified that at least one SV was significantly correlated with each of the known covariates (RIN, sequencing batch, PMI, race, and genotype-derived PCs). For each brain region, we corrected for their respective SVs and used residualized count matrices for differential expression analysis.

We calculated differential expression levels for each trauma measure (**Table 1**) in each brain region using limma^33^ with residualized expression for each gene as an outcome and trauma variable as a predictor. For sex-interactions, we used an interaction model with residualized expression for each gene as an outcome and an interaction term trauma variable:sex as a predictor including the main effects of trauma variable and sex in the model (Expression ∼ trauma x sex + trauma + sex). Differentially expressed genes (DEGs) and sex-interaction DEGs were defined by a nominal p-value cutoff of p <0.05 or an adjusted False Discovery Rate (FDR)<5%.

We next conducted a gene-level sensitivity analysis (Trauma Leading Gene Analysis), which identifies genes for which the variance explained in expression is greatest for trauma vs. sex or diagnosis. For each significant gene association, we determined which variables (PTSD, MDD, and/or sex) were associated with the gene in the same direction as trauma and, of those, which variable explained the most variance in the gene’s expression based on a nested R^2^ statistic in a joint model of expression (expression ∼ trauma + sex + diagnosis) (**SFig. 1A**).

We conducted gene set enrichment analysis of trauma-leading DEGs that were pre-ranked by association t-score using fgsea^34^ (R package). We tested for enrichment of a manually curated list of Gene Ontology terms relevant to postmortem neural tissue based on broad categories of terms (GO slims) accessed via QuickGO^35^. We then tested for overrepresentation of the functional categories among the significant (FDR<5%) GO terms using a binomial exact test.

We used Pearson correlation statistics of gene-trait t-scores to compare transcription profiles across brain regions as a genome-wide measure of concordance. In some cases we visualized the level of concordance/discordance using rank-rank hypergeometric overlap (RRHO) heatmaps where the input score was calculated as -log10(p-value)*sign(effect size) and all tested genes are included using RRHO2^36^(R package).

### Cell type-specific expression imputation and differential expression analysis

We deconvolved bulk brain tissue expression to estimate cell type proportions with single cell reference panels for cortex^37^ or amygdala^38^ using CIBERSORTx^39^. Excitatory and inhibitory neuronal subtypes were collapsed into single groups of excitatory neurons and inhibitory neurons. We tested for association of brain cell type proportions as an outcome and trauma variables, sex or diagnosis as a predictor using linear regression, covarying for sampling plate, postmortem interval, age, race and RIN. We then used bMIND^40^ to impute sample-level cell-type specific gene expression matrices for six brain cell types (excitatory neurons, inhibitory neurons, astrocytes, oligodendrocytes, endothelial cells, and microglia) for DLPFC, dACC, MeA and BLA, and an additional general “neurons” cell type for DLPFC and dACC.

As was done with bulk tissue, we calculated SVs for each cell type for each brain region and residualized cell type expression matrices for the optimum number of SVs as calculated in SVA. We ran differential expression analysis and trauma-leading gene analysis as previously described to obtain cell type DEGs (ctDEGs). FDR multiple testing correction was conducted within cell type within brain region with a 5% FDR significance threshold. Downstream analyses include only those ctDEGs which pass trauma-leading gene criteria.

To test whether some cell type was more responsive to one trauma variable category vs another, we performed a binomial test to assess whether the ratio of DEGs for each trauma type deviated from the proportion observed in bulk tissue samples for each cell type and brain region. P-values were adjusted for multiple comparisons using a FDR, with significance set at FDR<0.05. Adjusted log ratios were calculated by comparing the log ratio of DEGs in cell types to that in bulk tissue.

### Hierarchical clustering of transcriptional signatures

Transcriptional signatures were evaluated as gene-level Student’s T scores (logFC normalized by standard error) for genes nominally (p<0.05) significant in at least 1 trauma cdategory signature and included for hierarchical clustering based on Euclidian distances using hclust (method = “complete”). Clusters were then determined by cutting at inflection point of k-cluster Dunn index values (**SFig. 4C**) between 2 and 10 clusters. Trauma transcriptomic signature cluster- and brain region-associated genes were identified by association of gene-trauma t-scores with cluster identity or brain region vs all other clusters using linear regression.

### Gene Co-expression analysis

For coexpression analyses, we corrected gene expression count matrices for known covariates for each brain region^41^. Using variance partition, we identified factors that explain ≥1% variation in ≥10% of genes for each brain region and residualized out those effects. We then removed genes that are below the bottom 30^th^ percentile of median expression and dispersion (type= “cv”) of expression^42^. The resulting count matrix was used as input for coexpression module detection and module differential gene correlation analysis.

We analyzed gene co-expression using Multiscale Embedded Gene Co-expression Network Analysis (MEGENA) framework to construct planar filtered networks, cluster into multiscale modules and detect multiscale hubs for each brain region^43^. We used all samples available for each brain region to construct each multiscale coexpression network. Modules were required to have a minimum of 10 genes and a maximum of half of the genes detected for each brain region.

Modules were annotated for function and cell type based on module member enrichment of GO terms relevant to postmortem brain tissues and neural cell type-specific marker genes^44^ using the moduleGO function as part of the DGCA R package. The modules were then categorized into the function in which the top GO term belonged and the top cell type based on lowest significant p-value surviving a 5% FDR threshold. If a term was annotated to multiple categories, then the category representing the largest percentage of significant terms was assigned. We then tested for enrichment of trauma DEGs or sex-interacting trauma DEGs in modules using a Fischer’s Exact Test implemented in GeneOverlap R package. We also tested for overrepresentation of functional and cell type groups among trauma DEG-enriched modules using a binomial test where x=number of significant modules in a category X, n=total number of significantly modules and p=proportion of modules in category X of all modules. Binomial p-values<0.05 were considered nominally significant and FDR adjusted q-values<0.05 were considered significantly overrepresented.

Differential gene correlation analysis (DGCA)^45^ calculates a change in correlation between pairs of genes within a module for two conditions (e.g. trauma vs no trauma), which produces a scaled estimate (ZScoreDiff). We then calculated a median differential correlation (MeDC) value to obtain a module-level estimate of the change in coexpression structure with trauma exposure. For each module and trauma measure, we calculated MeDC values and corrected for multiple testing using a 5% FDR threshold for significance.

### Human induced pluripotent stem cell (hiPSC) transduction and treatment

Control hiPSC-derived NPCs (NSB553-S1-1, male, European ancestry; NSB2607-1-4, male, European ancestry) were cultured in hNPC media: DMEM/F12 (Life Technologies #10565), 1x N2 (Life Technologies #17502-048), 1x B27-RA (Life Technologies #12587-010), 1x Antibiotic-Antimycotic, 20 ng/ml FGF2 (Life Technologies) on Geltrex (ThermoFisher, A1413301). hiPSC-NPCs at full confluence (1-1.5×107 cells/well of a 6-well plate) were dissociated with Accutase (Innovative Cell Technologies) for 5 mins, spun down (5 mins X 1000g), resuspended and seeded onto Matrigel-coated plates at 3-5×106 cells/well. Media was replaced every two-to-three days for up to seven days until the next split.

At day −1, confluent hiPSC-NPCs were transduced with rtTA (Addgene 20342) and NGN2 (Addgene 99378) lentiviruses. At day 0 (D0), 1 µg/ml dox was added to induce NGN2-expression. At D1, transduced hiPSC-NPCs were treated with antibiotics to select for lentiviral integration (1 mg/ml G-418). At D3, NPC medium was switched to neuronal medium (Brainphys (Stemcell Technologies, #05790), 1x N2 (Life Technologies #17502-048), 1x B27-RA (Life Technologies #12587-010), 1 µg/ml Natural Mouse Laminin (Life Technologies), 20 ng/ml BDNF (Peprotech #450-02), 20 ng/ml GDNF (Peprotech #450-10), 500 µg/ml Dibutyryl cyclic-AMP (Sigma #D0627), 200 nM L-ascorbic acid (Sigma #A0278)) including 1 µg/ml Dox. 50% of the medium was replaced with fresh neuronal medium once every second day.

On day 5, young hiPSC-NPC NGN2-neurons were replated onto geltrex-coated plates. Cells were dissociated with Accutase (Innovative Cell Technologies) for 5-10 min, washed with DMEM, gently resuspended, counted and centrifuged at 1,000xg for 5 min. The pellet was resuspended in neuron media and cells were seeded at a density of 1.2×10^6^ per well of a 12-well plate.

At D11, iGLUTs were treated with 200 nM Ara-C (Sigma #C6645) to reduce the proliferation of non-neuronal cells in the culture, followed by half medium changes. At D17, Ara-C was completely withdrawn by full medium change. iGLUTs were fed by half medium changes until treated at D21 by a half medium change, with final concentrations of 1000nM HCort (Sigma #H0888), 10nM estradiol (Sigma #E1024), both or DMSO vehicle only. iGLUTs were harvested in trizol 48 hours post treatment.

### hiPSC RNA isolation, sequencing and analysis

Samples harvested in trizol were sent to the Yale Center for Genome Analysis for RNA isolation and library prep. Total RNA were run on the bioanalyzer for quality control, then selected for polyadenylated transcripts using oligo-dT beads followed by random priming. Sequencing was run on the Illumina NovaSeq instrument.

The Rsubread package^46^ was used to align raw reads to the GRCh38 genome assembly and to generate gene expression counts. Transcripts with at least 10 counts per gene in at least 1 sample were kept. Differential expression analyses were done using the limma package^33^, with voom-normalized counts and a design model including the drug treatment, technical replicate, RIN, RNA concentration, and surrogate variable identified by SVA. The limma::duplicateCorrelation function was applied in order to account for within-donor correlation. Genes with q_FDR_<0.05 were considered significantly differentially expressed.

## RESULTS

### Trauma is transcriptionally encoded in the human brain in a region- and cell-type-specific manner

To identify a transcriptional signature of trauma in the postmortem human brain, we first tested for transcriptional associations with trauma exposure (trauma_Any_) and cumulative trauma burden (trauma_Count_) as a quantitative measure across four brain regions: DLPLC, dACC, BLA and MeA. Our cohort includes 136 individuals with at least one reported trauma (168 had no reported trauma; **SFig. 1A**). Gene expression counts from RNA-seq were corrected for surrogate variables preserving the effects of trauma_Count_, sex and diagnosis and then tested for association with several trauma measures (**Table 1; STable 1**). We conducted sensitivity analyses to identify significant gene expression associations driven by trauma, rather than diagnosis or sex, termed trauma-leading genes, and only included those for downstream analyses (**See SMaterials; SFig. 1**).

We identified 4 genes whose expression significantly (p_FDR_<0.05) changed with increasing trauma_Count_ in the MeA: *LMO2* (logFC (per increase in 1 trauma exposure)= −0.034, p=2.2×10^−6^), *ARHGEF18* (logFC=0.010, p=4.1×10^−6^_)_, *ACVR1* (logFC=-0.021, p=4.2×10^−6^), and *SPTLC2* (logFC=0.014, p=7.92×10^−6^). 143-163 and 439-661 genes were nominally (p < 0.05) associated with trauma_Any_ and trauma_Count_, respectively, across each of the four brain regions (**Fig. 1A & SFig. 2A**). Using RRHO analysis, we observed a greater transcriptional concordance between prefrontal cortex regions vs amygdala regions among trauma_Count_ signatures (**Fig. 1B**). We conducted gene set enrichment analysis of all nominally significant (p<0.05) associations and enrichment of functional categories among those significantly enriched GO terms (**STable 2**). Trauma_Any_ signatures were enriched in gene regulation terms such as transcription factor binding (GO:0061629, NES=1.97, q_FDR_= 0.001) in the DLPFC, protein synthesis terms in dACC (GO:0003735, NES=2.71, q_FDR_= 3.8×10-11) and BLA (GO:1990904, NES=1.79, q_FDR_= 2.1×10-4) and spliceosome (GO:0005681 NES= 2.34, q_FDR_= 4.6×10-6) and mRNA processing (GO: 011441, NES=2.00, q_FDR_=4.6×10-6) in MeA. Trauma_Count_ signatures were enriched in immune response terms (downregulated; GO:0002274, NES=-2.2, q_FDR_=3.0×10-4) in DLPFC, and RNA processing (GO:0006396, NES=1.92, q_FDR_=4.25×10-8) and ion transmembrane transport (downregulated; GO:0015075, NES=-1.77, q_FDR_=1.43×10-4) in BLA (**Fig.1C-E**).

**Figure 1.**
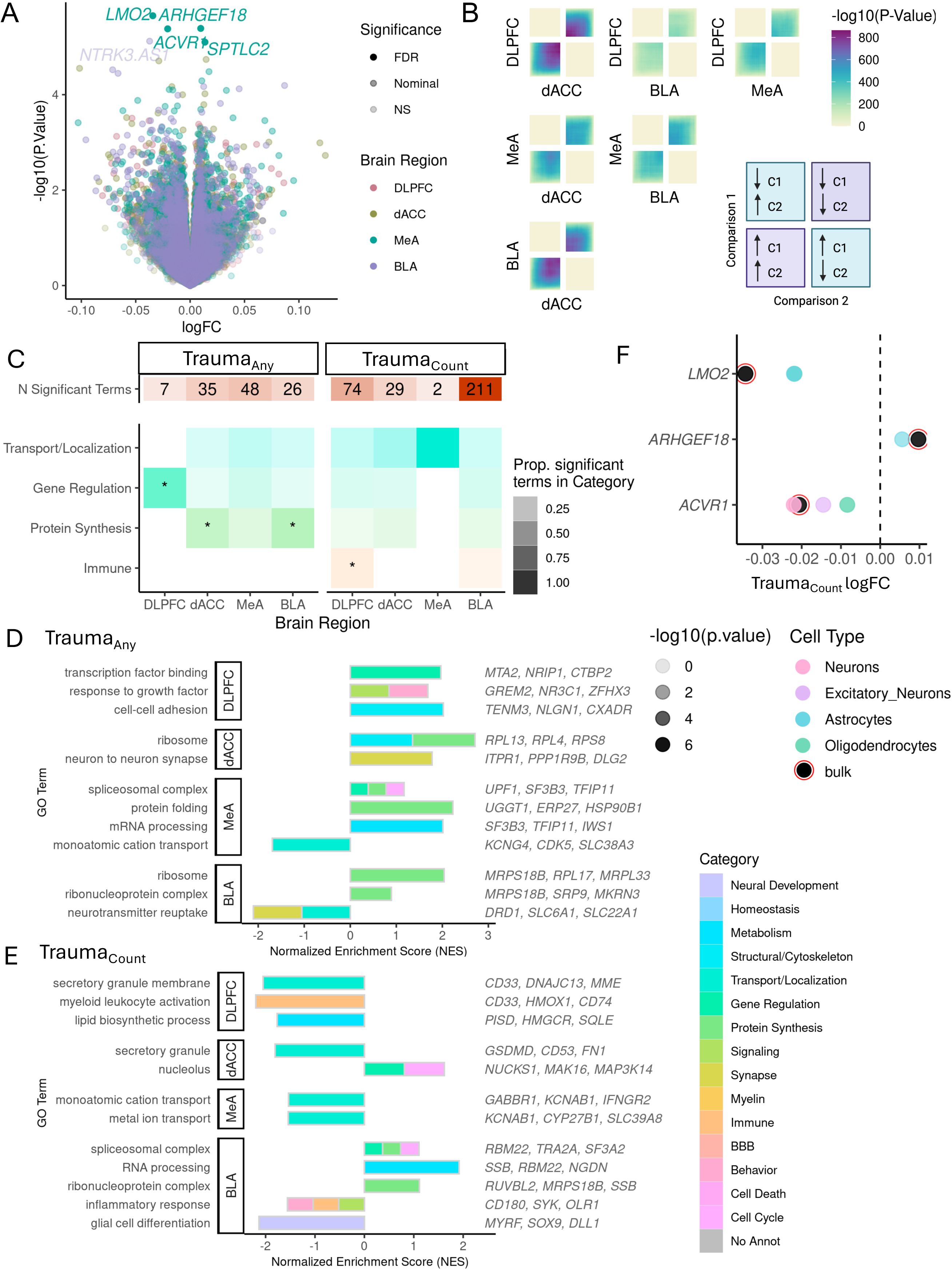
Transcriptional signatures of trauma exposure (Trauma_any_) and accumulation (Trauma_Count_). A. Gene-level expression associations with cumulative trauma (trauma_Count_). Log-fold-change (logFC) values associated with each additional trauma (1 unit increase in trauma load) and –log10 transformed raw p-values for each gene is shown. FDR significance < 5% and nominal significance is p< 0.05. Only trauma-leading gene associations are highlighted and included in follow up analyses. DLPFC = Dorsolateral prefrontal cortex, dACC=dorsal anterior cingulate cortex, MeA=Medial amygdala, BLA= basolateral amygdala. B. Concordance of trauma_Count_ transcriptional signatures for each brain region pair evaluated by stratified rank-rank hypergeometric overlap (RRHO) analysis. Bottom left quadrant indicates genes upregulated with trauma in both regions, upper right quadrant indicates genes downregulated with trauma in both regions. C. Gene Ontology term enrichment summary. Number of significant GO terms for each brain region for trauma_Any_ and trauma_Count._ Overrepresentation of significant GO terms by functional category. Shading of tile indicates proportion of significant GO terms in each category over the total number of significant terms for each brain region and trauma measure. Categories significantly overrepresented (q_FDR_ < 0.05) indicated with asterisk (*). D-E. Representative significant GO term associations from gene set enrichment analysis for trauma_Any_ (D) or trauma _Count_ (E) signature (p < 0.05 trauma-leading genes) for each brain region. Normalized enrichment score indicates degree of enrichment with positive direction indicating an enrichment of upregulated trauma genes and negative direction indicating an enrichment of downregulated trauma genes. Color of bar indicates functional category. F. Cell type differentially expressed genes (ctDEG) in association with trauma_Count_. Number of nominally significant (p<0.05) trauma-leading differentially expressed gene associations for each cell type (color bars) and brain region. G. MeA trauma_Count_ gene associations from bulk analyses are driven by specific cell types. Gene-level logFC estimates from bulk tissue expression (black with red circle) and from imputed cell type expression (color points). Only nominally significant (p<0.05) estimates shown.

Bulk gene expression analyses represent aggregate measures of cell type-specific expression. To isolate cell type-specific-patterns of trauma-gene associations, we deconvolved cell type proportions and imputed cell type gene expression using single cell RNA sequencing references from postmortem cortical and amygdala tissue ^37,38,40^. Cell type proportions across brain regions were significantly associated (p_FDR_ < 0.05) with sex and, for 5 brain region cell type-pairs, lowly correlated (r<|0.1|) with MDD diagnosis (**SFig. 2B&C**). We tested for associations of imputed cell-type expression with _traumaany_ and trauma_Count_ and identified 1-11 and 1-2091 cell-type DEGs (ctDEGs, p<0.001), respectively across regions(**STable 3**). Oligodendrocytes in dACC (2091 ctDEGs) and MeA (431 ctDEGs) had the largest transcriptional responses to trauma_Count_. We followed up on our significant MeA trauma_Count_ genes from bulk tissue (**Fig. 1A**) to identify potential cell types driving their association; *LMO2* (p=7.7×10^−6^) and *ARHGEF18* (p=2.1×10^−4^) associations were strongest in astrocytes and *ACVR1* trauma_Count_ association was strongest in neurons (p=1.×10^−6^) (**Fig. 1F**).

### Trauma disrupts gene coexpression and gene-gene correlations in a brain region-specific manner

To understand the systems-wide functional impacts of trauma, we used MEGENA to resolve 258, 208, 315, and 320 brain-region specific multiscale coexpression modules for DLPFC, dACC, MeA and BLA, respectively, and annotated each for functional and cell type enrichments (**SFig. 3; STable 4**). 41% (DLPFC), 37% (dACC), 41% (MeA) and 42% (BLA) of trauma signature genes were within coexpressed networks.. We hypothesized that these genes are coordinated by common regulators and carry out common functions making them ideal as potential therapeutic targets. We tested for enrichment of the trauma_any_ and trauma_Count_ transcriptional signatures in the corresponding brain region-specific co-expression modules (**Fig. 2A&B & SFig 4&5; STable 5**) and summarized the overrepresentation of neural functional and cell type categories among those enriched coexpressed modules (**Fig. 2C & SFig. 5C; STable 6**).

**Figure 2.**
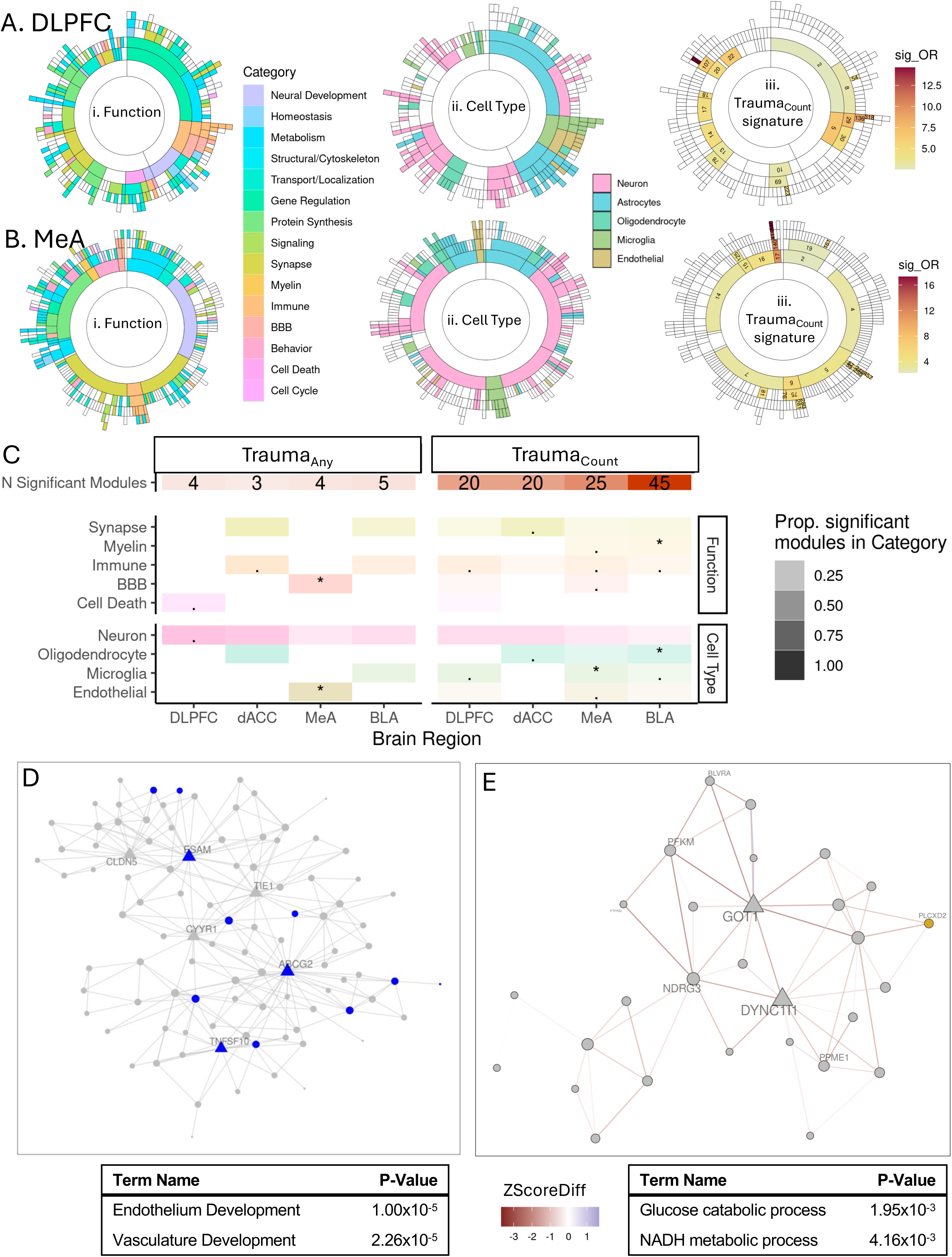
Trauma transcriptional signatures enrich in gene coexpression modules and alter coexpression relationships. A. Sunburst plot representing module hierarchy of DLPFC coexpression network where each arc represents a module. Functional category annotations (i), cell type annotations (ii) and enrichment odds ratio for modules significantly enriched (q_FDR_ < 0.05) for trauma_Count_ signature genes(iii). B. Sunburst plot representing module hierarchy of MeA coexpression network where each arc represents a module. Functional category annotations (i), cell type annotations (ii) and enrichments for traumaCount signature genes(iii) for each module. C. Module enrichment summary. Number of significantly enriched for each brain region for trauma_Any_ and trauma_Count._ Overrepresentation of significant modules by functional category. Transparency of tile indicates proportion of significant modules in each category over the total number of significant moduels for each brain region and trauma measure. Categories significantly overrepresented (q_FDR_ < 0.05) indicated with an asterisk (*). D. MeA module 140 enriched for MeA trauma_Count_ signature genes. Nodes represent module member genes and hubs genes are depicted as triangles. Genes downregulated with trauma_Count_ are shown in blue. Significant enriched GO terms of module and enrichment p-value shown. E. DLPFC module 224 is differentially correlated with trauma_Any_. Nodes represent module member genes and hubs genes are depicted as triangles. Edges indicate change in gene-gene-correlation (ZScoreDiff) between no trauma and trauma conditions. Negative ZScoreDiff indicates a decrease in correlation with trauma_Any_. Genes upregulated (yellow) and downregulated (blue) with trauma_Any_ are colored. Significant enriched GO terms of module and enrichment p-value shown.

Larger-scale modules were significantly enriched with trauma signature genes, and more modules were enriched in trauma_count_ signatures (20-45 modules across brain regions) than trauma_any_ (3-5 modules across brain regions). The BLA had the greatest number of modules enriched in the trauma_count_ signature with an overrepresentation of myelin (q_FDR_= 7.44×10^−3^) and oligodendrocyte (q_FDR_= 3.31×10^−3^) annotated modules (**Fig. 2C**). In the MeA, the dichotomous trauma_Any_ signature enriched in blood-brain barrier (BBB) (q_FDR_= 4.44×10^−3^) and endothelial modules (q_FDR_= 2.90×10^−2^) and the continuous trauma_count_ signature additionally enriched in microglia (q_FDR_= 0.046) modules. This suggests an overall disruption of MeA BBB function with exposure to any number of traumas and neuroimmune dysregulation with exposure to *increasing* traumas. Endothelial module 140 of the MeA was significantly enriched for both trauma_Any_ and trauma_Count_ signatures, specifically with genes downregulated with trauma_Count_ such as module hub genes *ABCG2* and *ESAM* (**Fig. 2D**).

While we may not see changes in a single gene’s expression following trauma, changes in coexpression module connectivity may suggest downstream dysregulation of their shared function. We tested for differential module connectivity, termed differentially correlated modules (DCMs), between samples with any trauma exposure and no reported trauma in each brain region (**SFig. 4; STable 7**) and tested for enrichment of functional and cell type categories (**SFig. 4E; STable 6**). Trauma_Any_ DCMs in the DLPFC were enriched for metabolic modules (binomial q_FDR_=0.049), such as module 224 annotated for neuronal metabolism (**Fig. 2E**). We found a gain in correlation for MeA module 499, which is annotated for lipid and sterol biosynthesis with hub gene *EPT1(* aka *SELENOI)* (**SFig. 4F**).

### Childhood trauma is characterized by amygdala and excitatory neuronal dysregulation

Lifetime traumatic experiences vary considerably on an individual level. The developmental period in which trauma occurs likely contributes to heterogeneity in transcriptional signatures of trauma. We repeated our differential expression and trauma driver analyses for samples from donors with trauma during childhood only (N=47) or adulthood only (N=45) compared to those with no reported trauma (N=162) (**STable 1**).Childhood trauma had a greater number of amygdala DEGs whereas adulthood trauma had a greater number of prefrontal cortical DEGs with a significant interaction between region and trauma (two-way ANOVA p=0.009) (**Fig. 3A**). Nominally significant (p<0.05) childhood trauma genes were enriched in ion transmembrane transport in MeA (downregulation; GO:0006812, NES=-1.73, q_FDR_=1.19×10^−4^) and BLA (GO:0015075, NES=-1.68, q_FDR_=2.79×10^−3^), and histone deacetylation in BLA (GO:0000118, NES=2.14, q_FDR_=8.05×10^−3^). Adulthood trauma genes were enriched in RNA processing and telomere lengthening in DLPFC (GO:0006397, NES=1.94, q_FDR_=1.62×10^−4^; GO:0010833, NES=2.13, q_FDR_=1.61×10^−3^) and MeA (GO:0006396, NES=1.81, q_FDR_=8.31×10^−6^) and ribosomal genes in DLPFC and BLA (**SFig.6A**; **STable 2**).

**Figure 3.**
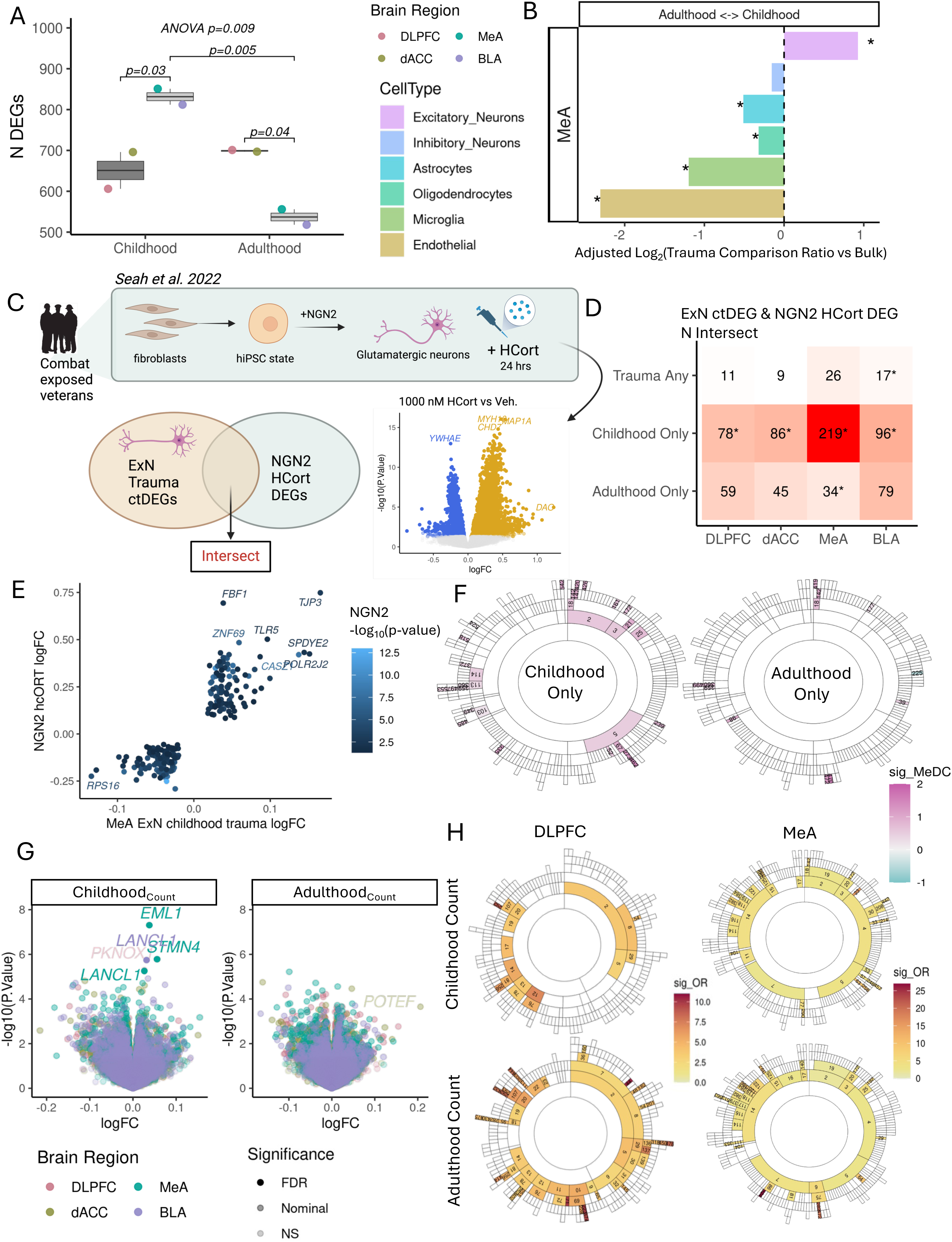
Childhood trauma and adulthood trauma have distinct transcriptional signatures. A. Number of childhood and adulthood DEGs (p <0.05) by brain region. Pvalues for two-way ANOVA and post-hoc Tukey for pairwise comparisons are annotated. DLPFC = Dorsolateral prefrontal cortex, dACC=dorsal anterior cingulate cortex, MeA=Medial amygdala, BLA= basolateral amygdala. B. MeA cell types are differentially responsive to childhood trauma compared to adulthood trauma. Log-transformed ratio of number of childhood ctDEGs over number of adulthood ctDEGs for each cell type (color bar) and adjusted for proportion of bulk childhood DEGs over bulk adulthood DEGs. Significant enrichment of number of childhood (positive value) or adulthood DEGS (negative value) by exact binomial test for deviation from bulk proportion with q_FDR_< 0.05 (*). C. Schematic of experiment applying 1000nM HCort to iPSCs-derived glutamatergic neurons and differentially expressed genes from Seah et al. 2022 and intersect of genes from Seah et al. 2022 and trauma ctDEGs from excitatory neurons (ExN) used in D. D. Intersect of genes from HCort DEGs (C) and ctDEGs from excitatory neurons (ExN) with respect to traumaany, childhood only and adulthood only across four brain regions. E. Concordantly regulated NGN2 HCort and MeA ExN childhood trauma genes by logFC. F. Childhood (left) and adulthood trauma (right) alter gene coexpression relationships among MeA coexpression modules. Significant differential correlated modules (DCMs; q_FDR_ < 0.05) are filled with module median differential connectivity score (MeDC) with negative values (cyan) indicating a loss of correlation among module member gene coexpression with trauma_Any_ and positive values (pink) indicating a gain of correlation.G. Gene-level expression associations with childhood trauma count (i) and adulthood trauma count (ii). Each point represents a gene, with the log-transformed fold change (logFC) per increase in 1 trauma on the x-axis and the -log10 transformed raw p-value on the y-axis. FDR significance < 5% and nominal significance is p< 0.05. Only trauma-leading gene associations are highlighted and included in follow up analyses. H. DLPFC and MeA modules and enrichment odds ratios for modules significantly enriched (q_FDR_ < 0.05) for childhood_Count_ and adulthood_Count_ signature genes.

Among the imputed cell type trauma DEGs (**SFig. 6B**), oligodendrocytes in dACC and inhibitory neurons in DLPFC had the greatest number of ctDEGs (p<0.05) in association with both childhood and adulthood trauma. We asked whether certain cell types were more transcriptionally responsive to childhood or adulthood trauma, as measured by a difference in the number of ctDEGs, than expected from bulk tissue DEG abundance. We observed the greatest differences in ctDEGs between childhood and adulthood trauma in the MeA: microglia and endothelial cells had more adulthood trauma ctDEGs, whereas excitatory neurons had more childhood trauma ctDEGs (**Fig. 3B**). Since limbic regions are not yet fully mature during childhood, we asked whether MeA excitatory neurons dysregulation with childhood trauma correlates with dysregulation observed in hiPSCs-derived NGN2 glutamatergic neurons exposed to HCort^28^(**Fig 3C**). As expected, childhood trauma ExN ctDEGs had the greatest overlap (219 concordant genes, Fisher’s exact p=3.2×10^−50^) with nominally significant HCort DEGs among excitatory neuron ctDEG signatures for trauma_Any_, childhood or adulthood traumas for each brain region (**Fig. 3D**). Concordantly regulated genes are shown in **Figure 3E**.

We tested for enrichment of childhood and adulthood trauma signatures in gene coexpression modules and for differential module connectivity following childhood or adulthood trauma (**SFig 7; STable 5&7)**. Among the significantly enriched modules, DLPFC modules annotated for neurons were overrepresented for both child and adult trauma signatures (**SFig. 6C; STable 6**). Though fewer MeA modules were significantly enriched in adult trauma, both childhood and adulthood trauma enriched modules shared subthreshold overrepresentations of neuron and BBB modules. Gene coexpression relationships were most disrupted in the MeA; 31 MeA modules were differentially correlated with childhood trauma and 12 MeA modules were differentially correlated with adulthood trauma with the majority of modules gaining connectivity with trauma (**Fig. 3F**). We did not observe any significant enrichments of functional or cell type categories among MeA DCMs (**SFig. 6D**).

We hypothesized that some genes may be associated with the accumulation of childhood or adulthood traumas, which may not be captured in the binary variables (Childhood only vs no trauma and adulthood only vs no trauma). We repeated analyses with a continuous variable for cumulative childhood(childhood _count_) or adulthood traumas(adulthood_count_), and dentified 4 significant childhood_count_ DEGs (q_FDR_<0.05) in the MeA (*AIRN*, logFC=-0.11, p=6.2×10^−9^; *EML1*, logFC=0.04, p=5.0×10^−8^; *STMN4*, logFC=0.06, p=1.7×10^−6^; and *LANCL1*, logFC=0.03, p=5.6×10^−6^), one of which (*LANCL1*, logFC=0.03, p=1.8×10^−6^) was also significantly associated in the BLA (**Fig. 3G; STable 1**).

At a transcriptome-wide scale, we observed brain region specificity in the response to childhood_count_ compared to adulthood_count_. Of all nominally significant DEGs (p<0.05), the frequency of DEGs across brain regions remained similar to trauma_Count_, with the exception of nearly double the genes in the DLPFC associated with adulthood_count_ (**SFig. 8A**). Childhood_count_ signatures were enriched in cis-golgi (downregulation) in DLPFC, ion transport (downregulation) in MeA and DNA repair in BLA; adulthood_count_ signatures were enriched in mRNA processing in the DLPFC and mitochondrial and protein synthesis genes (**SFig. 8B; STable 2**). Transcriptome-wide correlations across signatures and hierarchical clustering also demonstrated a divergence in the encoding of increasing child and adult traumas. While the childhood_count_ signatures largely correlated and clustered with trauma_Count_, they negatively correlated with (r=-0.18 to r=-0.28) and clustered away from adulthood_count_ (**SFig. 8C-D**).

Among coexpression modules, adulthood_count_ signatures were enriched in 60 DLPFC modules with an overrepresentation of neuronal modules (**Fig. 3H & SFig. 9A; STable 6**). This is in contrast to the 17 DLPFC modules enriched in childhood_count_ signatures with an overrepresentation of protein synthesis modules, supporting a strong encoding of adulthood trauma in the DLPFC. In the MeA, childhood_count_ and adulthood_count_ signatures were largely enriched in similar modules (**Fig. 3H**), with similar subthreshold overrepresentations in oligodendrocyte and myelin modules (**SFig. 9B**). However, we identified a significant enrichment of immune/microglia modules (MeA module 6 and child modules 75, 282, 472) in adulthood_count_ signatures but not enriched with childhood_count_ signatures.

### Transcriptomic signatures of trauma differ by trauma type

Transcriptional differences between childhood and adulthood trauma may be a result of differences in the types of traumas children and adult experience. To address this, we repeated our differential expression and downstream analyses (**SFig. 10A-C; STable 2**) and compared the transcriptional signatures of developmental age (childhood, adulthood, or both) with that of different types of traumas (combat, interpersonal, or both; single acute, complex, or both) and trauma load (high or low trauma). We also dissected phenotypic relationships among trauma variables in this cohort with multiple correspondence analysis (MCA) on donor trauma profiles alone (**Table 1-2**, **Fig. 4A, SFig. 11**). Dimension 1 of MCA explains the correspondence of trauma exposed categories from the ‘no trauma’ categories. Dimensions 2 and 3 reveal several clusters of corresponding trauma variable categories for which we have assigned the following identifiers (**Fig. 4B**): (1) <u>High Load</u>: *both childhood and adulthood traumas*, *both combat and interpersonal traumas, both single and complex traumas* and *top quartile of trauma* categories; (2) <u>Complex</u>: *interpersonal only*, *childhood only* and *complex only*; (3) <u>Adulthood</u>: *adulthood onl*y and *bottom quartile;* (4) <u>Combat</u>: *combat trauma only* and (5) <u>Single</u>: *single acute trauma only*.

**Figure 4.**
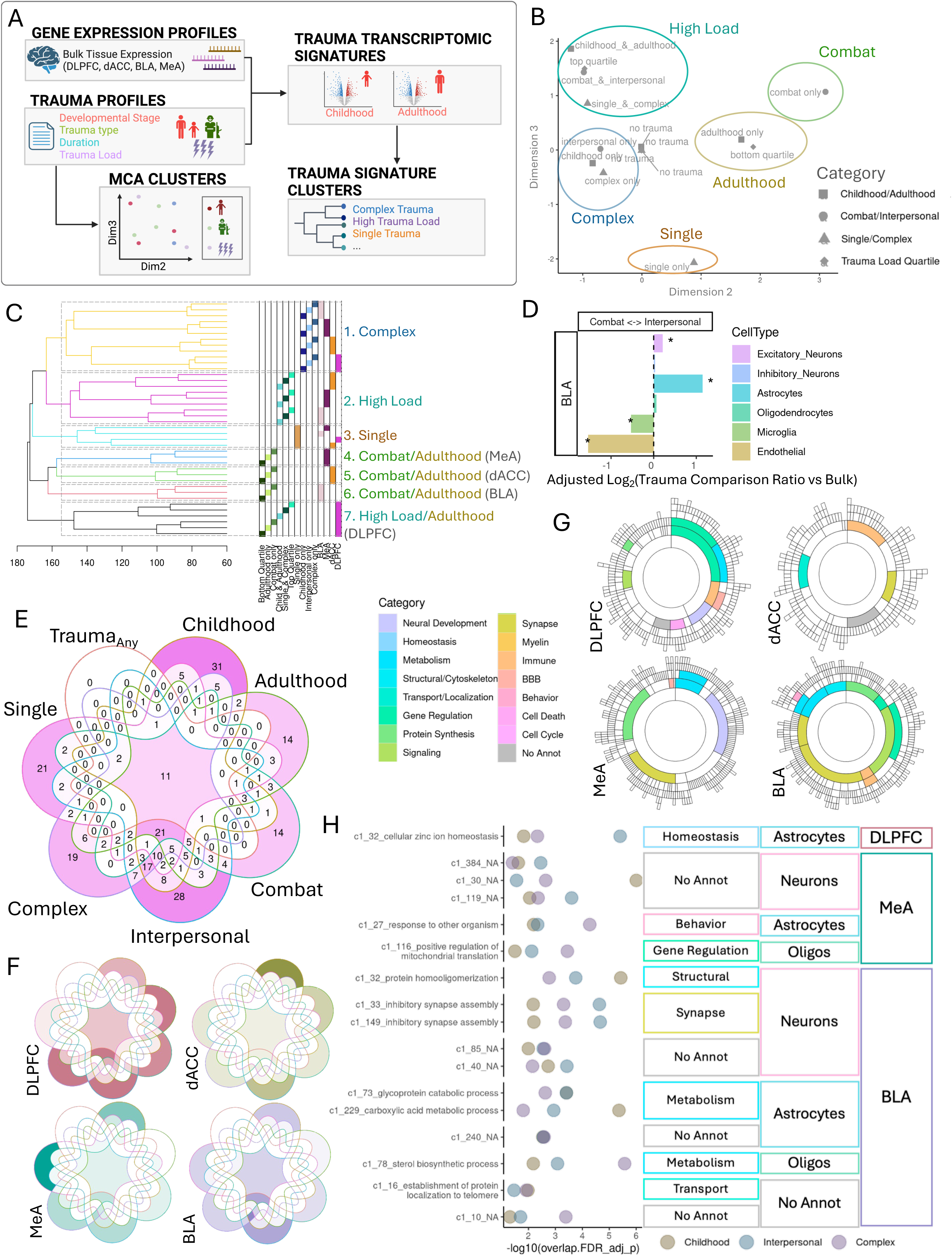
Trauma types have shared and distinct transcriptional signatures A. Analysis of trauma variable profiles and their transcriptomic signatures. B. Multiple correspondence analysis (MCA) of donor trauma profiles. Dimensions 2 and 3 reveal clusters of similar trauma variable categories, which we categorize into five MCA clusters: High load, complex traumas, adulthood, single and combat. C. Hierarchical clustering of trauma variable category transcriptional signatures. Clustering heights are shown on the x axis. Trauma variable and brain region for each transcriptomic signature is annotated to the right by color and in text with congruent MCA clustering categories. Dashed rectangles designate trauma transcriptional signature clusters. D. Log-transformed ratio of number of interpersonal ctDEGs over number of combat ctDEGs for each cell type (color bar) and adjusted for proportion of bulk interpersonal DEGs over bulk combat DEGs. Significant enrichment of number of interpersonal (positive value) or combat DEGS (negative value) by exact binomial test for deviation from bulk proportion with q_FDR_< 0.05 (*). E-F. Intersections of the number of modules enriched for each trauma type signature for all brain regions combined (E) and separated by brain region (F). G. Modules commonly enriched for all 6 trauma types (childhood, adulthood, combat, interpersonal, complex and single trauma) and colored by annotated functional category. H. Modules commonly enriched for childhood, interpersonal and complex trauma signatures. Module enrichment log-transformed FDR-adjusted p-value shown for each trauma signature (color). Brain region, cell type and functional annotations shown. Modules are labeled with significant (q_FDR_< 0.05) GO term with the lowest pvalue.

We then asked whether trauma category transcriptomic signatures were more similar within MCA trauma category clusters or within brain regions using a hierarchical clustering approach (**Fig.4C**). Based on clustering of transcriptomic associations alone, we identified 7 clusters of trauma type-transcriptional signatures. Clustering revealed patterns of region-specific responses to different dimensions of trauma. In general, MCA trauma profile clusters remained clustered together and separate from other trauma profile clusters at greater heights (h=162.5-171.9); combat transcriptional signatures tended to cluster with adulthood transcriptional signatures. Within MCA trauma profile clusters, transcriptional signatures clustered by brain region (h=126.2-162.4), and then by individual trauma variable categories (h=75.5-112.2). Combat/adulthood traumas signatures demonstrated the greatest transcriptional differences between brain regions compared to other trauma categories. There was one exception to this general pattern: DLPFC transcriptional signatures for adulthood and high load trauma category clusters separated away from dACC and amygdala signatures for those trauma categories (h=174.1). This suggests that exposure to adulthood trauma is encoded similarly in DLPFC regardless of past traumas in childhood.

Using our imputed cell type expression, we asked whether certain cell types were more transcriptionally responsive to specific trauma types (**SFig. 10D**) and differentially responsive between trauma variable categories (**SFig. 10E**). While most ctDEGs across brain regions were comparatively enriched for interpersonal traumas, BLA endothelial cells and to a lesser extent, BLA microglial, dACC neurons and MeA oligodendrocytes were more responsive in combat trauma profiles **(Fig. 4D).**

We next looked at coexpression modules enriched in trauma type transcriptional signatures (**STable 7**) and their overlap across combat, interpersonal, single, complex, childhood and adulthood, as well as trauma_Any_. Overall, the majority of modules were either specific to one trauma or enriched for all trauma types, with the exception of a cluster of common modules between interpersonal, complex and childhood traumas. No modules were specific to Trauma_Any_ (**Fig. 4E**), which suggests that these 6 trauma types reflect the heterogeneity in trauma signatures among coexpressed genes. 32 modules in total (10, 4, 6, and 12 modules in DLPFC, dACC, MeA and BLA), were commonly enriched among all trauma types (**Fig. 4E&F**). These convergent modules were spread across all 4 brain regions and various functions and cell types (**Fig. 4G, SFig.12**). Of those modules specific to one trauma type, childhood trauma-specific modules were mostly in dACC (14 modules) (**SFig.11, SFig.14A**), adulthood-specific modules were split between cortical regions (DLPFC 7, dACC 6), combat-specific modules were greatest in the DLPFC (6 modules)(**SFig.14B**), interpersonal-specific modules were split among all brain regions (5(MeA)-8(BLA&dACC)), complex-specific modules were greatest in DLPFC and single trauma-specific modules were greatest in MeA (13 modules) (**Fig. 4F**; **SFig. 12**). 17 modules were commonly enriched among childhood, interpersonal and complex trauma types: 11 of those were in the BLA annotated with neuronal and metabolic functions, for example, inhibitory synapse assembly (modules 33 and 149) in neurons, and glycoprotein (module 73), carboxylic acid metabolism (module 229) in astrocytes and sterol biosynthesis (module 78) in oligodendrocytes. (**Fig.4H**).

### Convergent sex specific mechanisms of trauma and HCort

Previous studies have demonstrated sex-specific transcriptional alteration with stress-related psychiatric disorders (see Introduction). To identify genes for which the effects of trauma are modulated by sex, we tested for trauma x sex interaction on expression for genes that were nominally significant trauma DEGs (**Fig 5A**). Of the 25,876 gene-trauma associations tested, 9 associations survived FDR correction and 405 associations (345 unique genes) were nominally significant for trauma-by-sex interaction (**SFig. 15A, STable 9)**, including *SLC38A3*(interaction beta=-0.18145785, p=0.0003882087) and *PPARD* (interaction beta=-0.03932737, p=0.0002782924; **Fig. 5B&C**). To determine whether the 9 significant trauma x sex interactions were driven by specific cell types, we tested their association within imputed cell type expression (**SFig. 15B**). We identified 7 genes (*SLC38A3, PPARD, SNORA46, MIR4292, IL1RN, DLX5, PCNA*) for which neurons had the strongest trauma x sex interaction, including *SLC38A3* (interaction beta=-0.22, p=0.0002) and *PPARD* (interaction beta= −0.032, p=0.0063) (**Fig. 5D**). We tested for enrichment of nominally significant sex-interacting trauma genes in brain region-specific coexpression modules and found 30 module-trauma variable enrichments (p_FDR_<0.05) across the four brain regions studied (**STable 8**). Several modules were significantly enriched for genes relatively upregulated in females with interpersonal traumas in dACC and childhood trauma in MeA (**Fig. 5E)**. In the dACC, module 368 was enriched for female-specific childhood trauma and interpersonal trauma genes (interpersonal: OR=537.1, p=1.26×10^−5^, q_FDR_= 0.003) and was annotated for astrocytes and with hub genes *TLR4*, *ETNPPL* and *P2RY1* (**Fig. 5F)**. In the MeA, module 6 was enriched for genes upregulated specifically in females (OR=69.4 p=2.8×10^−6^, q_FDR_=8.7×10^−4^) such as *HCG22*, *CLEC7A*, *ARHGAP15* and *TMC8* and was annotated for immune response and microglia (**SFig. 15C**).

**Figure 5.**
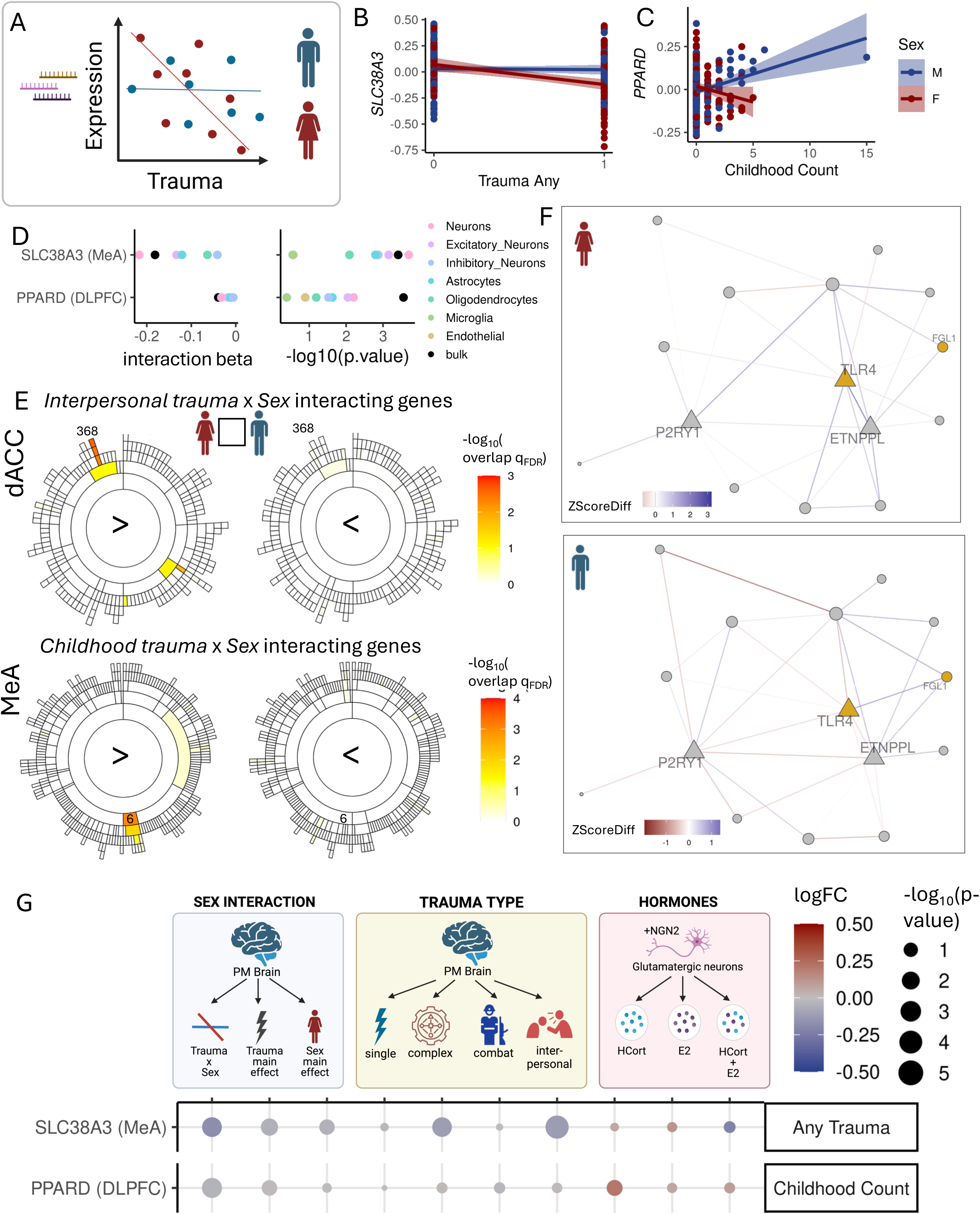
Sex moderates the impact of trauma on the transcriptome and gene-coexpression networks. A. Interaction effect of sex and trauma on gene expression. B. SLC38A3 expression (residualized) with trauma exposure in females (red) and males (blue). C. DLX5 expression (residualized) in males (blue) and females (red) with single only, complex only, both single & complex trauma or no trauma. D. Sex-interaction term estimates (left) and log transformed pvalues (right) for SLC38A3 with trauma exposure and DLX5 with single trauma from bulk tissue expression (black with red circle) and from imputed cell type expression (colored points). E. Enrichment pvalues of sex-interactive interpersonal trauma genes (top) or childhood trauma genes (bottom) with relative upregulated in females (left) vs relative upregulation in males (right) in dACC (top) or MeA (bottom) multi-scale coexpression modules. F. Coexpression network graph of dACC module 368. Nodes are colored by significant sex-interacting genes (yellow = upregulate in females) and edges are colored by the SE normalized difference in the correlation (ZscoreDiff) interpersonal trauma only to no trauma condition in females (top) and in males (bottom). ZscoreDiff>0 indicates a gain in correlation in expression of the two node genes with trauma and ZscoreDiff<0 indicates a loss of correlation. G. Gene expression association estimates for 2 significant sex-interacting genes (y-axis) in various contexts (x-axis). From this dataset: sex-interaction term estimate with trauma variable category (righthand box), main effect of trauma variable category, main effect of sex, main effect of single trauma, complex trauma, combat trauma, and interpersonal trauma. From RNA seq of human NGN2 neurons derived from induced pluripotent stem cells with in vitro exposure to hydrocortisone (HCort), estradiol (E2) or a combination of both (HCort+E2). Point color indicates the logFC value and size indicates –log10 transformed pvalues of those estimates.

We next explored potential mechanisms for sexual dimorphism in gene expression with trauma. We asked whether these associations are driven by sex-biases in types of trauma exposure, by exposures to sex-specific hormones or their interaction with stress hormones. To directly test hormonal effects in a cell type-specific manner, we applied HCort, estradiol (E2), and their combination (HCort+E2) to hiPSC-derived NGN2 glutamatergic neurons *in vitro* and identified 12, 0 and 39 significant (q_FDR_<0.05) DEGs, and 787, 677, 642 nominally significant (p<0.01) DEGs respectively, for each condition (**SFig16A&B**).

For significant (q_FDR_ < 0.05) trauma x sex-interacting genes, we compared gene expression associations with different trauma types within this postmortem dataset and with HCort, E2 and HCort+E2 exposure in the NGN2 neuronal cell populations (**Fig.5B,SFig. 16C**). We highlight 2 genes: *SLC38A3* and *PPARD*. Expression of *SLC38A3* in the MeA is specifically decreased in females with trauma exposure (**Fig. 5B**). sWe observed a similar downregulation of *SLC38A3* with interpersonal traumas (logFC=-0.12, p=5.6×10^−6^), suggesting this sex-interaction may be due to an increased prevalence of interpersonal traumas in females (**Fig 5F**). *PPARD* expression in DLPFC was specifically downregulated in females and upregulated in males with increasing childhood_count_(**Fig. 5C**), with the childhood_count_ x sex interaction driven by neurons (**Fig 5D**). We observed a similar upregulation in iPSC-derived NGN2 neurons treated with only HCort (logFC=0.21, p=0.02), but not when combined with estradiol (HCort+E2; logFC=0.1, p=0.3), which suggests a hormonal mechanism for the observed sex interaction with childhood_count_.

## DISCUSSION

We investigated the transcriptional signatures of trauma exposure (trauma_any_), trauma accumulation (trauma_count_), traumas at distinct developmental periods (childhood only, adulthood only, childhood_count,_ and adulthood_count_), distinct trauma types (combat, interpersonal, complex, and single traumas), and trauma load (high load and low load) across four corticolimbic regions of the postmortem human brain. Using imputed brain-region and cell type-specific signatures, cell type annotation of coexpression modules and hiPSC-derived neuronal populations, we observed dysregulation of each neural cell type and each brain region to at least some aspect of trauma exposure, accumulation or trauma type. Lastly, we tested for sex-specificity of trauma transcriptional regulation and identified potential mechanisms (trauma type and hormonal) driving sexual dimorphism in transcription of each sex-specific trauma gene.

We used a hierarchical clustering approach to compare transcriptional signatures of various trauma types. Generally, we found that transcriptional signatures are more variable between trauma measures than between brain regions. DLPFC signatures remained the most similar across all trauma measures that included an exposure to adulthood trauma. This suggests that the DLPFC remains plastic in its encoding of traumas in adulthood regardless of past childhood trauma experiences. On the other hand, we saw the greatest transcriptional differences between brain regions (MeA, BLA and dACC) for adulthood and combat traumas and greater transcriptional similarity in amygdala and dACC with childhood traumas. Together, this evidence suggests that traumas occurring earlier in life have similar widespread changes in gene expression in the amygdala, but as the frontal cortex and amygdala mature, transcriptional effects of trauma diverge. In other words, the amygdala may encode earlier trauma that persists through the lifespan, whereas the prefrontal cortex remains transcriptionally flexible in response to additional traumas.

This supports current evidence for amygdala involvement in childhood trauma^47^ and weakened functional connectivity between prefrontal cortex and amygdala as a result of early trauma^48–50^. Rodent models of early life stress tell a similar story of persistent alteration in amygdala circuitry and function and anxiety-like behaviors even after the stress is removed^51–53^, with evidence for the MeA in particular being involved in mediating sex differences in responses to early stress ^54^.

We investigated transcriptional responses of specific neural cell types using imputed cell type expression, providing insights into lasting functional effects at the cellular level. In both cortical regions, we observed elevated inhibitory neuron transcription across several trauma measures (aside from combat trauma with an increased endothelial response in the BLA), which supports the current evidence of gamma-aminobutyric acid (GABA)-ergic dysregulation in chronic stress, depression and PTSD ^26,55,56^. We similarly found a large transcriptional response of oligodendrocytes particularly in the dACC with most trauma types. Oligodendrocytes and myelination have also previously been implicated in MDD ^57^, childhood abuse^58^ and stress pathophysiology^59^, with impaired myelination induced in mouse prefrontal cortex in response to chronic social stress^60^

We also observed a relatively larger response of MeA excitatory neuron transcription to childhood trauma compared to adulthood traumas (**Fig.3B**). Using a hiPSC-derived neuronal model, we identified genes concordantly regulated by childhood trauma and by HCort, a glucocorticoid responsible for systemic recruitment of stress response. Excitatory neurons within the MeA and its circuitry are not yet fully mature during childhood when the trauma occurs; likewise, hiPSCs-derived neurons modeled *in vitro* are thought to retain an immature state^61^. The identified concordant genes may reflect persistent transcriptional alterations specific to the GC signaling as a result of childhood trauma. Thus, postmortem brain tissues and hiPSC-derived neural cells make complementary systems to dissect molecular mechanisms of neuropsychiatric disorder risk.

Using gene coexpression network analysis, we further investigated the enrichment of trauma DEGs in multi-scale modules of coexpressed genes and, orthogonally, the impact of trauma in dissolving or strengthening module coregulatory relationships. Across all trauma measures, we observed an enrichment of large-scale modules; generally, gene regulation and synaptic neuronal modules in cortical regions and neuronal, glial and immune modules in amygdala. However, as modules specialize, enrichment of trauma DEGs diverge to different small-scale modules by different trauma types. With our differential gene correlation analyses, we found that coregulatory relationships of MeA modules were most altered by trauma, supporting our differential expression results, though trauma DEG enriched modules and differentially correlated modules were, generally, minimally overlapping. These trauma-associated modules and their central hub genes are ripe for further dissection for dissection of their relationship with mood and anxiety symptoms.

Our study highlights the importance of considering different types of trauma, such as those during distinct developmental periods (i.e., childhood trauma versus adulthood trauma) and trauma type (complex, interpersonal, combat and single acute traumas). While analyses that consider all trauma aggregations yield significant power, we also demonstrate as well that distinct trauma types have distinct transcriptomic impacts that affect different pathways and cell types. Among coexpressed networks of genes, all trauma_Any_ signature enriched modules were shared with at least one trauma type module, suggesting a robust dissection of the heterogeneity encompassed within the trauma_Any_ signature by trauma types. We have previously demonstrated considerable heterogeneity in the genetic predisposition to PTSD based on trauma type, specifically between a military vs civilian cohort^16^. This approach is increasingly important as we disentangle the heterogeneity of psychiatric diagnoses^62^.

As stated earlier, postmortem studies have identified substantial sex differences of brain gene expression in PTSD and MDD (see Introduction); and on a population scale, there are sex biases in the prevalence diagnosis of psychiatric diagnoses ^5^ and in their comorbidities^64^. Here, we investigated sex-specific transcriptional effects of trauma and found several sex x childhood trauma interactions in MeA, supporting the involvement of MeA in mediating sex differences in response to early stress in rodent studies^54^.

Since sex is multifaceted and interrelated with gender, we consequently investigated a number of biological and societal factors that may drive sex differences in transcription. First, we considered the role of trauma type and influences of gendered social experiences. We identified solute carrier *SLC38A3* (*SNAT3*), which is involved in glutamate-GABA-glutamine cycling, as being downregulated in females with trauma exposure (trauma_Any_), but observed a similar downregulation with respect to interpersonal trauma. Women are more likely to experience (and report) sexual traumas whereas men are more likely to experience combat and assault^65–67^. Even when accounting for trauma type, women are still more likely than men to develop PTSD following sexual trauma ^67^. Further, levels of social and societal support and their potential to mediate psychopathology differ by trauma type and gender^68,69^.

We also considered the impacts of estradiol, alone and in concert with hydrocortisone on human induced NGN2 neurons to assess possible contributions of sex hormones in transcriptional responses to stress^19,70^. The *PPARD* transcript encodes a peroxisome proliferator-activated receptor (PPAR) which is highly expressed in neurons, astrocytes and glia and linked to the development of anxiety^71^. DLPFC expression of *PPARD* was specifically upregulated in males with accumulating childhood trauma compared to that in females. Similarly, this transcript was upregulated with treatment of HCort, but not in the additional presence of E2 in hiPSC-derived NGN2 neurons suggesting a hormonal basis of sex interaction with childhood trauma at a neuronal level. Together, we resolved not only trauma-signatures in a sex-specific manner, but also pinpointed aspects of biological sex which may drive these differences for a subset of our trauma-associated genes.

As with all postmortem studies, identifying transcriptomic signatures of trauma or other exposures requires careful delineation of cause and effect; signatures identified may represent the impact of predisposition to the exposure, diagnosis relating to the exposure, medications taken to alleviate symptoms, or the exposure itself. In the case of trauma, genes identified may represent predisposition to PTSD/MDD, or to trauma itself. For example, volunteering for military service has a genetic component^72,73^. They may also be associated with diagnoses stemming from trauma, or may reflect medications taken to mitigate symptoms stemming from trauma. We apply two approaches to mitigate this concern. First, we test for PTSD and MDD signatures, and note that these are distinct from (although correlated with) our trauma signatures. However, it is possible that these case/control analyses are underpowered compared to our quantitative trauma analysis, and that analysis of PTSD or MDD severity or symptom domains (e.g., see^26^) may yield additional gene associations. Second, we apply sensitivity analyses to retain only those gene-trauma associations for which trauma explains the most variance in expression compared to diagnosis (or sex). As such, we are left with fewer genes overall, but those for which we have the highest statistical confidence for association with trauma.

This study, to our knowledge, presents the largest postmortem brain transcriptomic study of psychosocial trauma. We approach trauma as a quantitative trait and dive into the heterogeneity of traumatic events, disentangling common and divergent molecular mechanisms of developmental period and trauma type. Future efforts in brain donation should focus on obtaining trauma histories for all individuals, especially those with no psychiatric diagnoses despite lifetime trauma, to study stress resilience and to reduce confounding factors in postmortem brain transcriptomic studies.

## Supporting information

SFig1

SFig

SFig3

SFig4

SFig5

SFig6

SFig7

SFig8

SFig9

SFig10

SFig11

SFig12

SFig13

SFig14

SFig15

SFig16

**SFigure 1.** Trauma Leading Gene Analysis. A. Frequency of trauma exposure (trauma any; top) and cumulative trauma (trauma count; bottom) and filled by Sex (left) or Diagnosis (right). B. Trauma, sex, gender and diagnosis are interrelated in this cohort. For each gene significantly associated (p<0.05) with some trauma variable, we ask whether trauma explains the most amount of variance in gene expression in a joint model using a nested R^2 for variables (Sex and/or diagnosis) with the same direction of effect as trauma. C. Gene-level estimates (logFC; top) and -log10 transformed pvalues (bottom) for all genes significant in at least one of 4 analyses: trauma exposure, cumulative trauma, PTSD case status and MDD case status. D. Genome-wide correlations of logFC values (top) or pvalues (bottom) for all genes nominally significant in at least one of the two transcriptional signature being compared. E. Breakdown of all genes nominally significantly associated with trauma exposure (trauma_any) or cumulative trauma (trauma_count) by driver variable from trauma driver analysis (A). F. Examples of gene expression ∼ trauma associations that are Trauma-, Sex-, PTSD-, and MDD-Leading. Each point represents the nested R^2 values from a joint model (see A) for each variable in the direction of effect on gene expression (sign(logFC)).

**SFigure 2.**Transcriptional signature of trauma_any_ and cell type deconvolution. A. Gene-level differential expression associations with trauma (trauma_Any_) compared to no trauma. Each point represents a gene, with the log-transformed fold change (logFC) on the x-axis and the - log10 transformed raw p-value on the y-axis. FDR significance < 5% and nominal significance is p<0.05. Only trauma-leading gene associations are highlighted and included in follow up analyses. DLPFC = Dorsolateral prefrontal cortex, dACC=dorsal anterior cingulate cortex, MeA=Medial amygdala, BLA= basolateral amygdala. B. Proportions of cell types from cell type deconvolution of bulk RNA sequencing for each brain region. C. Association between cell type proportion by brain region with trauma variables, diagnosis and sex. Asterisks annotates significant associations (q_FDR_ < 0.05). D. Cell type differentially expressed genes (ctDEG) in association with trauma_Any_. Number of nominally significant (p<0.05) trauma-leading differentially expressed gene associations for each cell type (color bars) and brain region.

**SFigure 3.** MEGENA coexpression modules and annotations. Brain-region specific multiscale coexpression modules for DLPFC, dACC, MeA and BLA resolved using MEGENA. Sunburst plot representing module hierarchy of coexpression networks where each arc represents a module. Each module is annotated for a neural functional category (left) and cell type category (right) based on gene set overlap (q_FDR_ < 0.05) with GO terms and neural cell type gene sets.

**SFigure 4.** Coexpression module enrichment and differential correlation with trauma exposure (Trauma_Any_). A-D. Sunburst plot representing module hierarchy of coexpression network where each arc represents a module. Functional category annotations (top left), cell type annotations (top right). Trauma_Any_ transcriptional signatures enrichments in brain region-specific coexpression modules (bottom left). Enrichment odds ratios of significantly enriched (q_FDR_ < 0.05) modules are highlighted from yellow to red. Trauma_Any_ alters gene coexpression relationships among coexpression modules (bottom right). Significant differential correlated modules (DCMs; q_FDR_ < 0.05) are filled with module median differential connectivity score (MeDC) with negative values (cyan) indicating a loss of correlation among module member gene coexpression with trauma_Any_ and positive values (pink) indicating a gain of correlation. E. Number of trauma_Any_ DCMs (orange-red) and functional and cell type category enrichment of trauma_Any_ DCMs by brain region. Shading of tile indicates proportion of significant modules in each category over the total number of significant modules for each brain region. Categories significantly overrepresented (q_FDR_ < 0.05) indicated with asterisk (*) or nominally significant with a period(.). F-G. MeA module 499 is differentially correlated with trauma_Any_. Nodes represent module member genes and hubs genes are depicted as triangles. Edges indicate change in gene-gene-correlation (ZScoreDiff) between no trauma and trauma conditions. Negative ZScoreDiff indicates a decrease in correlation with trauma_Any_. Genes upregulated (yellow) and downregulated (blue) with trauma_Any_ are colored. Significant enriched GO terms of module and enrichment p-value shown.

**SFigure 5.** Enrichment of trauma transcriptional signatures in coexpression modules. A-B: Sunburst plot representing module hierarchy of dACC(A) and BLA (B) coexpression network where each arc represents a module. Functional category annotations (i), cell type annotations (ii) and enrichments for traumaCount signature genes(iii) for each module. Enrichment odds ratios of significantly enriched (q_FDR_ < 0.05) modules are filled from yellow to red. C. Enrichment of trauma transcriptional signatures in brain region coexpression modules and their functional and cell type categories. The total number of coexpression modules (red orange) enriched for trauma_Any_ and Trauma_Count_ signatures for each brain region. The proportion of significant modules within a functional or cell type category over the total number of significant modules. An asterisk (*) indicates those categories overrepresented among trauma signature enriched categories based on exact binomial test (q_FDR_ < 0.05) and dot indicates nominal significance (p < 0.05). BBB=blood-brain barrier

**SFigure 6.** Enrichments of transcriptional signatures for childhood trauma or adulthood trauma among bulk, tissue cell types and coexpression modules. A. Gene Ontology term enrichment summary. Number of significant GO terms for each brain region for childhood only and adulthood only. Overrepresentation of significant GO terms by functional category. Shading of tile indicates proportion of significant GO terms in each category over the total number of significant terms for each brain region and trauma measure. Categories significantly overrepresented (q_FDR_ < 0.05) indicated with asterisk (*). Enriched GO term associations from gene set enrichment analysis of childhood only or adulthood only transcriptional signatures (p < 0.05 trauma-leading genes) for each brain region. Normalized enrichment score indicates degree of enrichment with positive direction indicating an enrichment of upregulated trauma genes and negative direction indicating an enrichment of downregulated trauma genes. Color of bar indicates functional category.B. Top:Cell type differentially expressed genes (ctDEG) in association with trauma_Count_. Number of nominally significant (p<0.05) trauma-leading differentially expressed gene associations for each cell type (color bars) and brain region. Bottom: Comparing the number of childhood ctDEGs to the number of adulthood ctDEGs for each brain region and cell type. Log-transformed ratio of number of childhood ctDEGs over number of adulthood ctDEGs for each cell type (color bar) and adjusted for proportion of bulk childhood DEGs over bulk childhood DEGs. Significant enrichment of number of childhood (positive value) or adulthood DEGS (negative value) by exact binomial test for deviation from bulk proportion with q_FDR_< 0.05 (*). C. Number of modules for each brain region significantly enriched for childhood only and adulthood only transcriptional signatures. Overrepresentation of significant modules by functional category. Shading of tile indicates proportion of significant modules in each category over the total number of significant modules for each brain region and trauma measure. Categories significantly overrepresented (q_FDR_ < 0.05) indicated with asterisk (*). D. Number of modules differentially correlated (DCMs) with childhood and adulthood trauma for each brain region. Overrepresentation of significant DCMs by functional category. Shading of tile indicates proportion of significant modules in each category over the total number of significant modules for each brain region and trauma measure. Categories significantly overrepresented (q_FDR_ < 0.05) indicated with asterisk (*).

**SFigure 7.** Modules differentially correlated with childhood trauma or adulthood trauma. Sunburst plot representing module hierarchy of coexpression network where each arc represents a module. Childhood only (upper) and adulthood only (lower) transcriptional signatures enrichments in brain region-specific coexpression modules (left). Odds ratios of significantly enriched (q_FDR_ < 0.05) modules are highlighted from yellow to red. Significant differential correlated modules (right; DCMs; q_FDR_ < 0.05) are filled by module median differential connectivity score (MeDC) for each respective trauma and brain regsion; negative values (cyan) indicating a loss of correlation among module member gene coexpression with trauma and positive values (pink) indicating a gain of correlation.

**SFigure 8.** Transcriptomic signatures for accumulation of childhood or adulthood traumas. A. Number of childhood_Count_ and adulthood_Count_ DEGs (p <0.05) by brain region. Pvalues for two-sample t-test was used to compare the number of DEGs between adulthood and childhood in amygdala and cortex. DLPFC = Dorsolateral prefrontal cortex, dACC=dorsal anterior cingulate cortex, MeA=Medial amygdala, BLA= basolateral amygdala. B. i.Gene Ontology term enrichment summary. Number of significant GO terms for each brain region for childhood count and adulthood count transcriptional signatures. Overrepresentation of significant GO terms by functional category. Shading of tile indicates proportion of significant GO terms in each category over the total number of significant terms for each brain region and trauma measure. Categories significantly overrepresented (q_FDR_ < 0.05) indicated with an asterisk (*). Enriched GO term associations from gene set enrichment analysis of childhood only (ii) or adulthood only(iii) transcriptional signatures (p < 0.05 trauma-leading genes) for each brain region. Normalized enrichment score indicates degree of enrichment with positive direction indicating an enrichment of upregulated trauma genes and negative direction indicating an enrichment of downregulated trauma genes. Color of bar indicates functional category. C. Childhood count and adulthood count transcriptional signatures are negatively correlated. Pairwise transcriptome-wide correlations (r) of trauma-gene logFC values for all genes nominally significant in at least one of the two transcriptional signatures being compared for each pair. D. Hierarchical clustering dendrogram of brain region-specic childhood count and adulthood count transcriptional signatures.

**SFigure 9.** Coexpression enrichments of transcriptomic signatures for accumulation of childhood or adulthood traumas. A. Sunburst plot representing module hierarchy of dACC and BLA coexpression network where each arc represents a module. Functional category annotations (i), cell type annotations (ii) and enrichments for traumaCount signature genes(iii) for each module. Enrichment odds ratios of significantly enriched (q_FDR_ < 0.05) modules are filled from yellow to red.

B. Number of modules for each brain region significantly enriched for childhood count and adulthood count transcriptional signatures. Overrepresentation of significant modules by functional category. Shading of tile indicates proportion of significant modules in each category over the total number of significant modules for each brain region and trauma measure. Categories significantly overrepresented (q_FDR_ < 0.05) indicated with asterisk (*).

**SFigure 10.** Trauma type transcriptomic signatures. A-B. Number of significant (q_FDR_< 0.05 ;A) and nominally significant (p < 0.05; B) DEGs for each trauma variable category and brain region. C. Gene Ontology term enrichment summary. Number of significant GO terms for each brain region for trauma type signautres. Overrepresentation of significant GO terms by functional category. Shading of tile indicates proportion of significant GO terms in each category over the total number of significant terms for each brain region and trauma measure. Categories significantly overrepresented (q_FDR_ < 0.05) indicated with asterisk (*). D. Cell type differentially expressed genes (ctDEG) for trauma type signatures and brain region. Number of nominally significant (p<0.05) trauma-leading differentially expressed gene associations for each cell type (color bars). E. Comparing number of ctDEGs for opposing trauma types: childhood vs. adulthood, interpersonal vs. combat and complex vs. single trauams. Log-transformed ratio of number of ctDEGs for each trauma type pair (color bar), adjusted for the same proportion from bulk tissue. Significant deviation (q_FDR_< 0.05 *) from bulk ratio tested for each cell type by exact binomial test.

**SFigure 11.** MCA analysis of donor trauma profiles A. Scree plot of MCA dimensions and proportion of explained variances B. Loadings of each trauma variable category on dimensions 1-5. C-D. MCA plot dimensions 1 vs 2 (C) and dimensions 2 vs 3 (D) of each of the variable categories with points labeled by cos2.

**SFigure 12.** Intersections of the number of modules enriched for each trauma type signature by brain region.

**SFigure 13.** Modules commonly enriched for all 6 trauma type signatures. Module enrichment log-transformed FDR-adjusted p-value shown for each trauma signature (color). Brain region, cell type and functional annotations shown. Modules are labeled with significant (q_FDR_< 0.05) GO term with the lowest pvalue.

**SFigure 14.** Modules only significantly enriched for childhood (top) or combat (bottom) trauma signatures. Module enrichment log-transformed FDR-adjusted p-value shown for each trauma signature (color). Brain region, cell type and functional annotations shown. Modules are labeled with significant (q_FDR_< 0.05) GO term with the lowest pvalue.

**SFigure 15.** Genes whose expression is significantly moderated by sex. A. Significant sex interaction term estimates for gene (y axis) with trauma variable category (box to the right). Point size indicates log transformed pvalues and color indicates which sex the gene is comparatively upregulated in. B. Sex-interaction term estimates (left) and log transformed pvalues (right) for significant sex interacting genes with trauma variable category (righthand box) from bulk tissue expression (black with red circle) and from imputed cell type expression (colored points). C. Coexpression network for MeA module 6 with sex differential genes with interpersonal trauma highlighted; yellow indicates upregulated in females and blue indicates upregulate in males. Edges are colored by the SE normalized difference in the correlation (ZscoreDiff) from interpersonal trauma only to no trauma condition in females (left) and in males (right). ZscoreDiff >0 indicates a gain in correlation in expression of the two node genes with trauma and ZscoreDiff < 0 indicates a loss of correlation.

**SFigure 16.** iPSC derived neuronal expression with hCORT and expression of sex-interacting genes across different trauma types postmortem and experimental conditions *in vitro*. A. Schematic of iPSC-induced glutamatergic neurons and application of HCort, E2 and HCort+E2 and RNA sequencing. B. Volcano plots of differentially expressed genes of NGN2 glutamatergic neurons from A. C. Gene expression association estimates for significant sex-interacting genes (y-axis) in various contexts (x-axis). From this dataset: sex-interaction term estimate with trauma variable category (righthand box), main effect of trauma variable category, main effect of sex, main effect of single trauma, complex trauma, both single&complex trauma, combat trauma, and interpersonal trauma. From RNA seq of human NGN2 neurons derived from induced pluripotent stem cells with in vitro exposure to hcort, estradiol or a combination of both. Point color indicates the logFC value and size indicates – log10 transformed pvalues of those estimates.

## ACKNOWLEDGEMENTS

We thank Rebecca Signer, Ada Kepinska, Hannah Young and Kayla Retallick-Townsley for critical feedback on the manuscript. We are grateful to the families who donated to this research.

