## Supplementary figures and images for "Decoding the transcriptomic signatures of psychological trauma in human cortex and amygdala"

### SFig

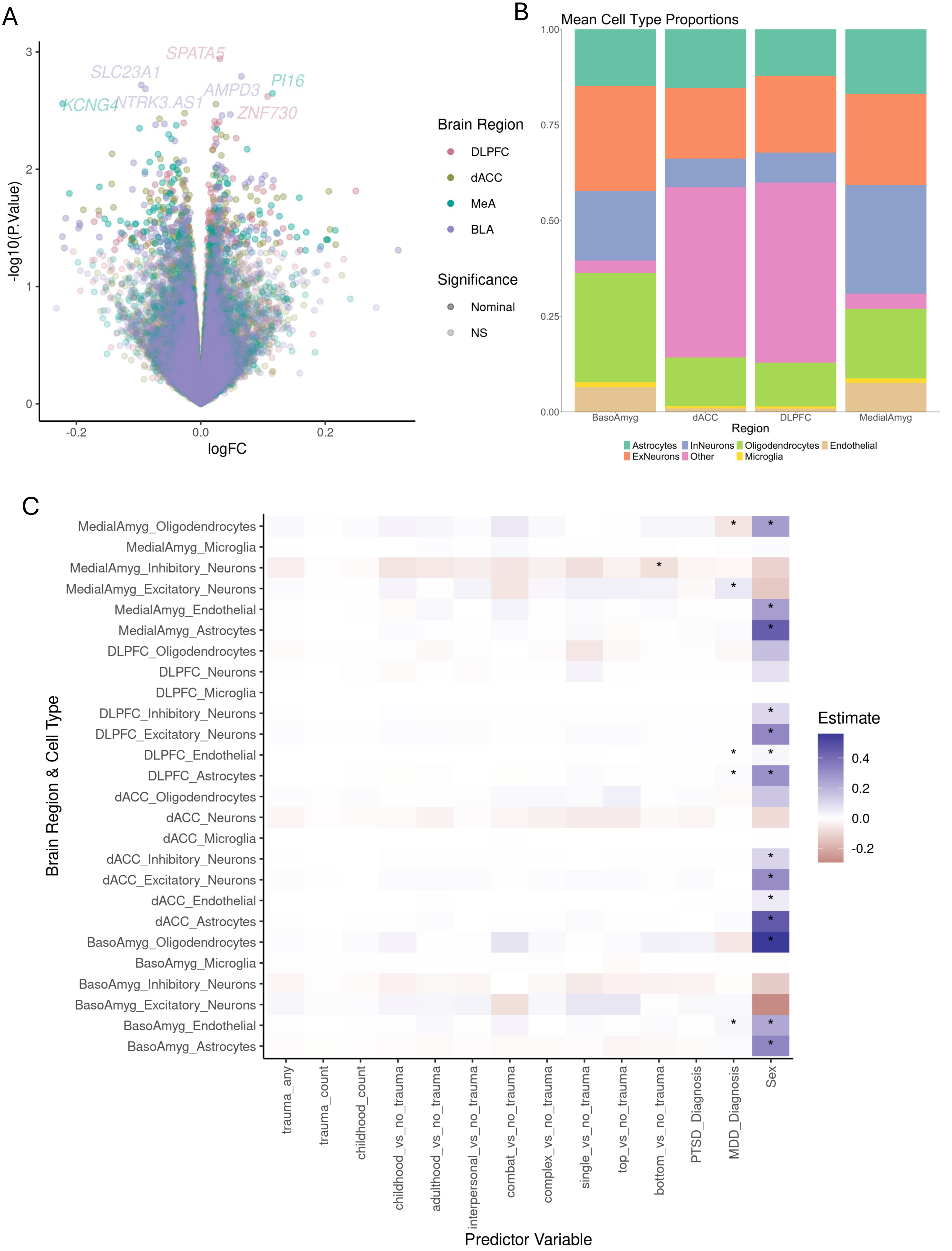

### SFig1

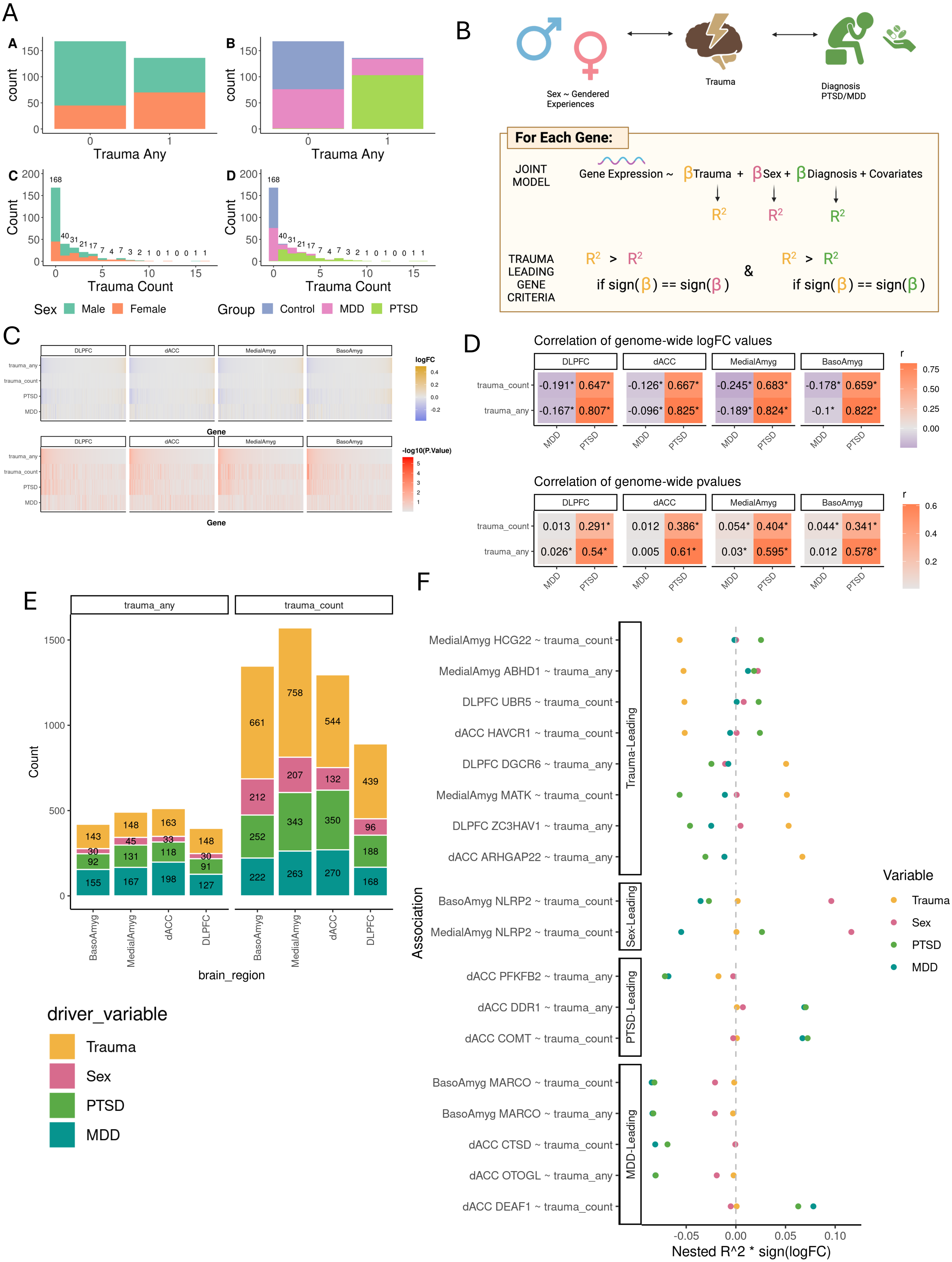

### SFig3

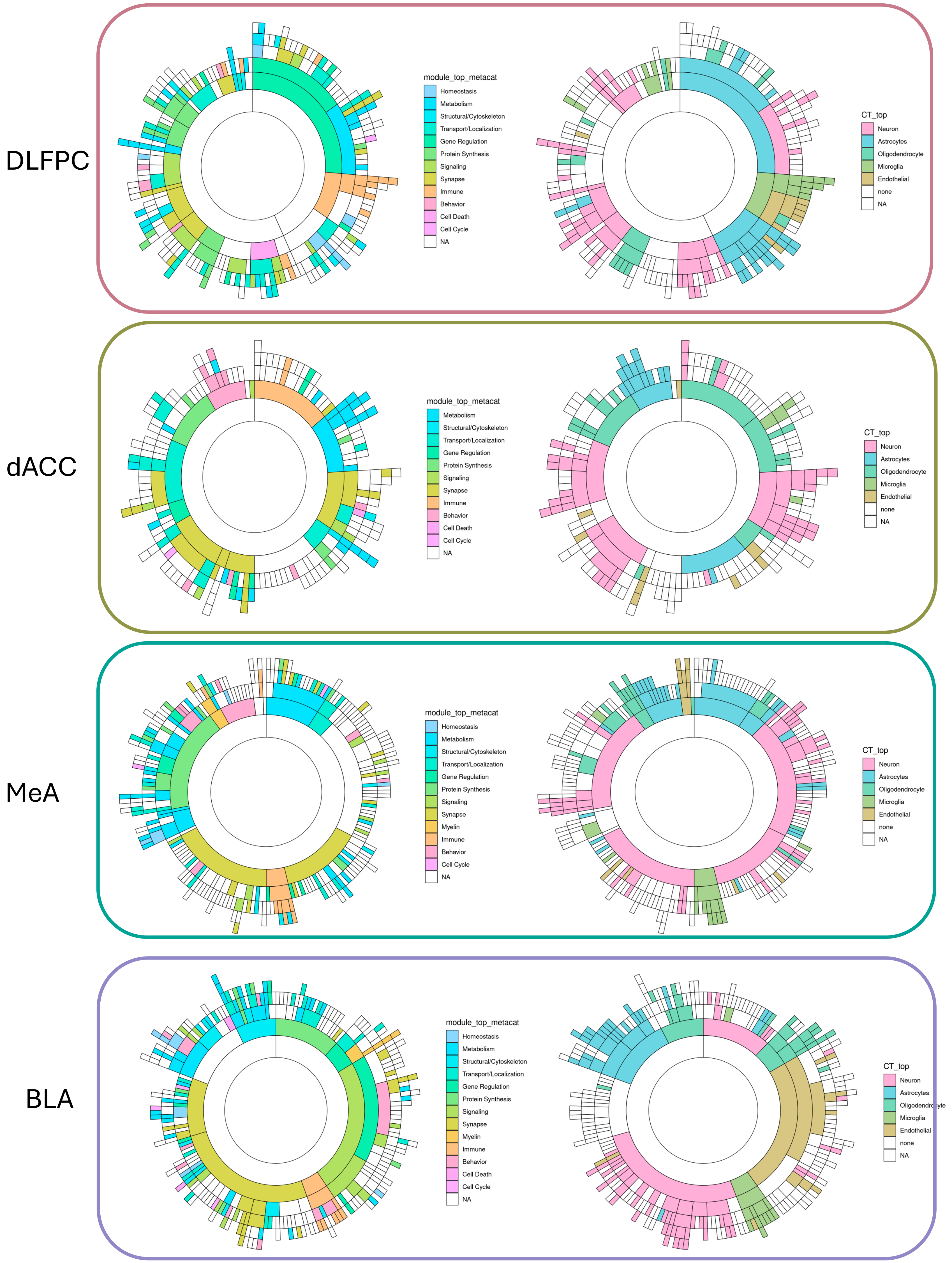

### SFig4

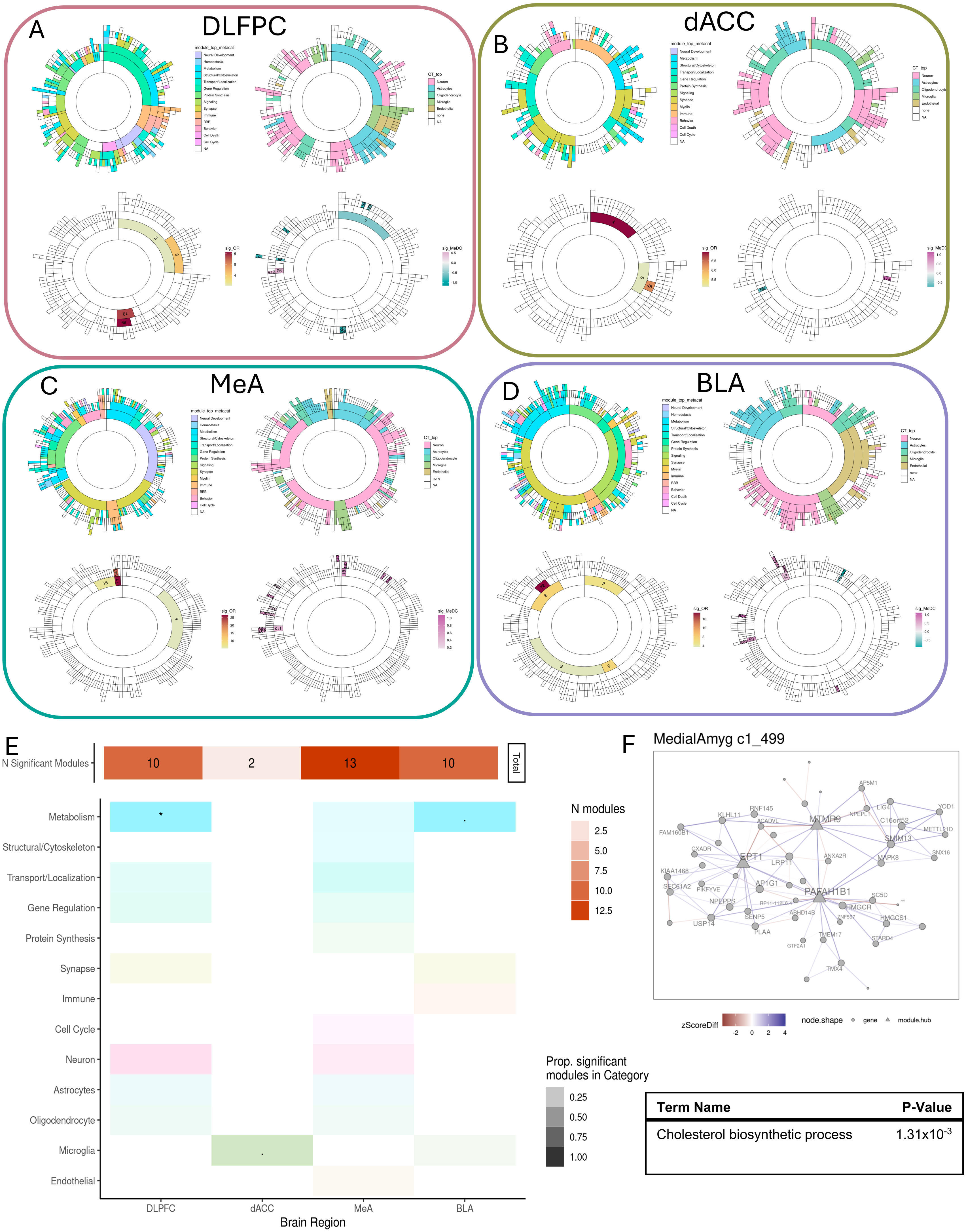


| Term Name | P-Value |
|---|---|
| Cholesterol biosynthetic process | $1.31x10^{-3}$ |

### SFig5

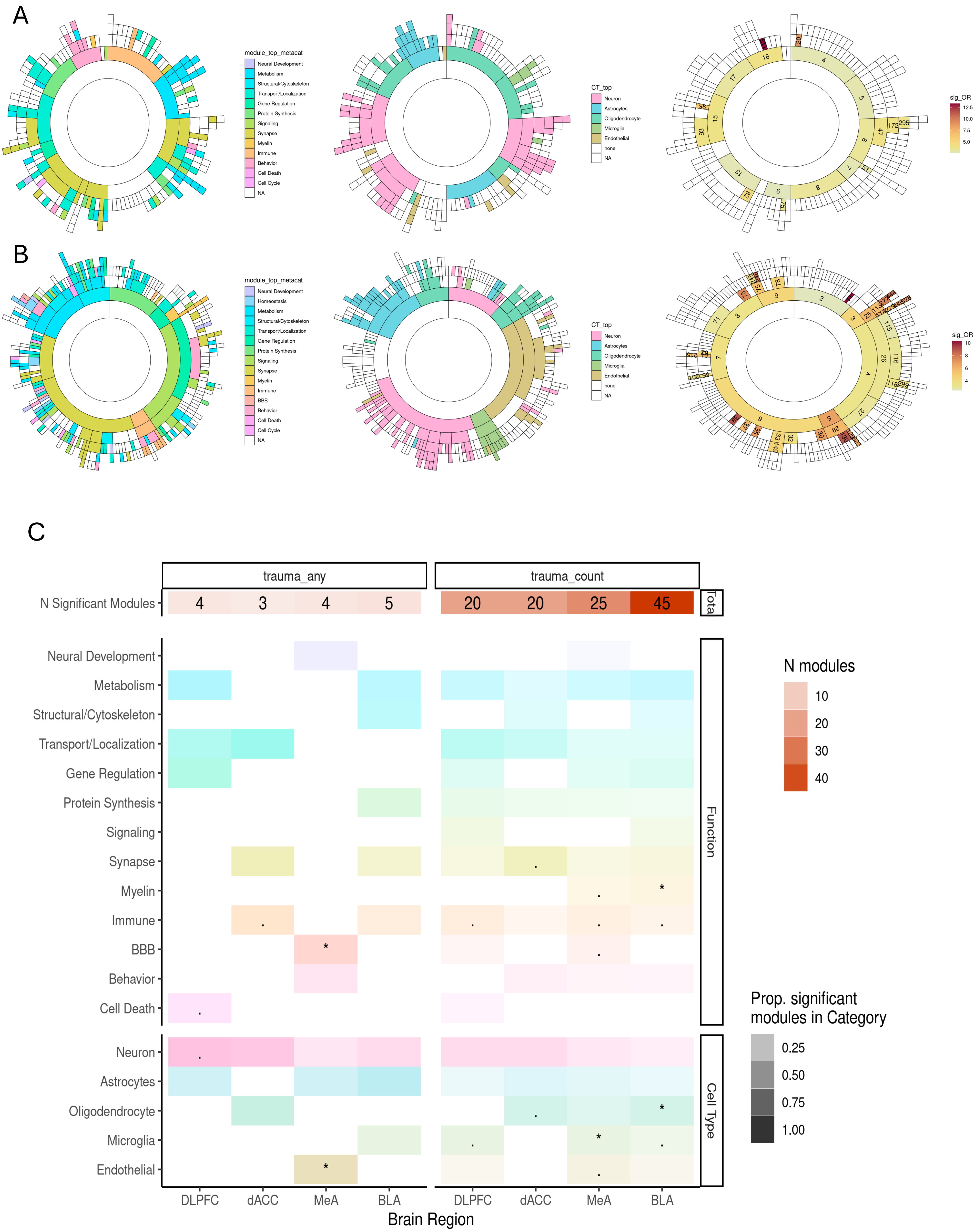

### SFig6

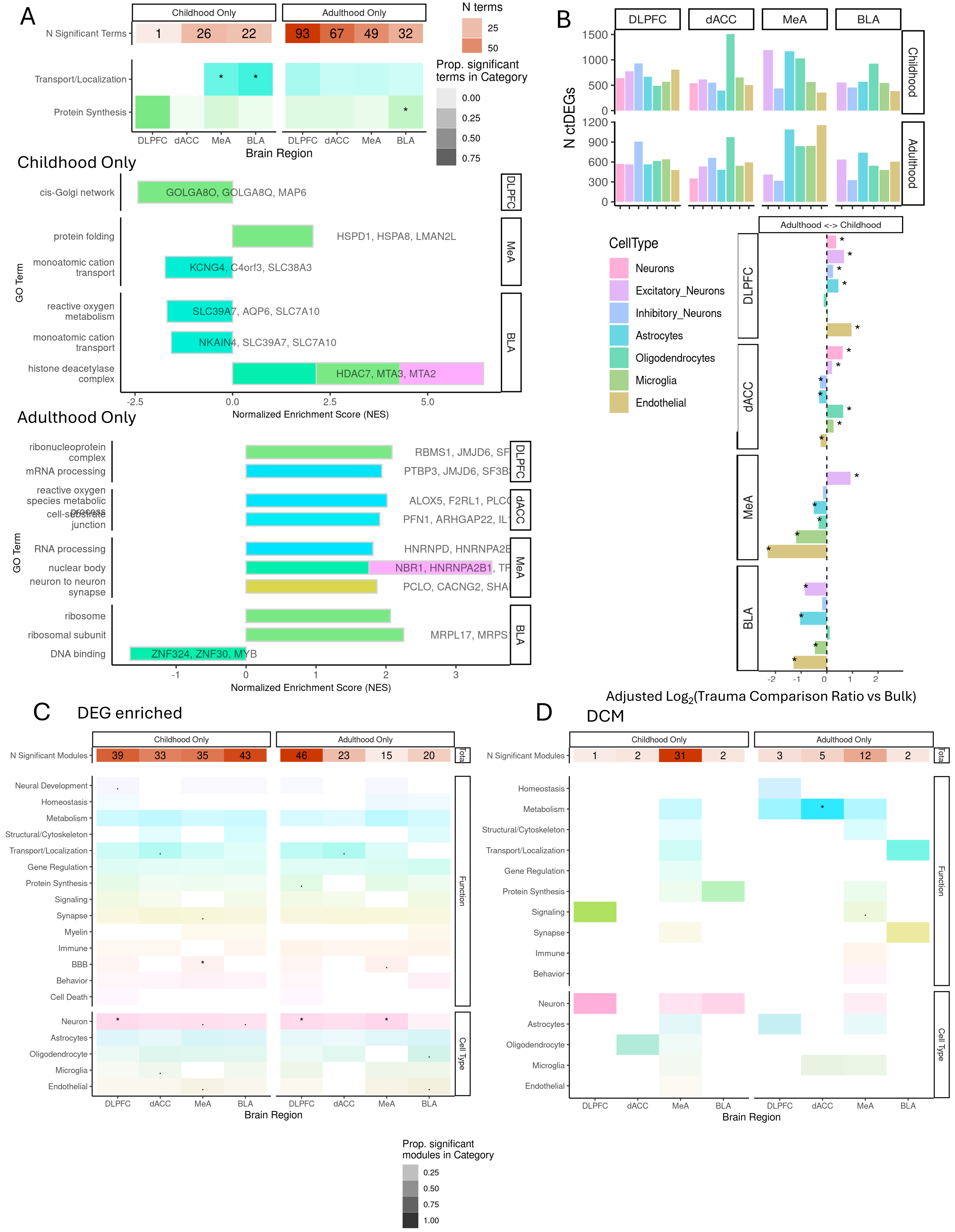

### SFig7

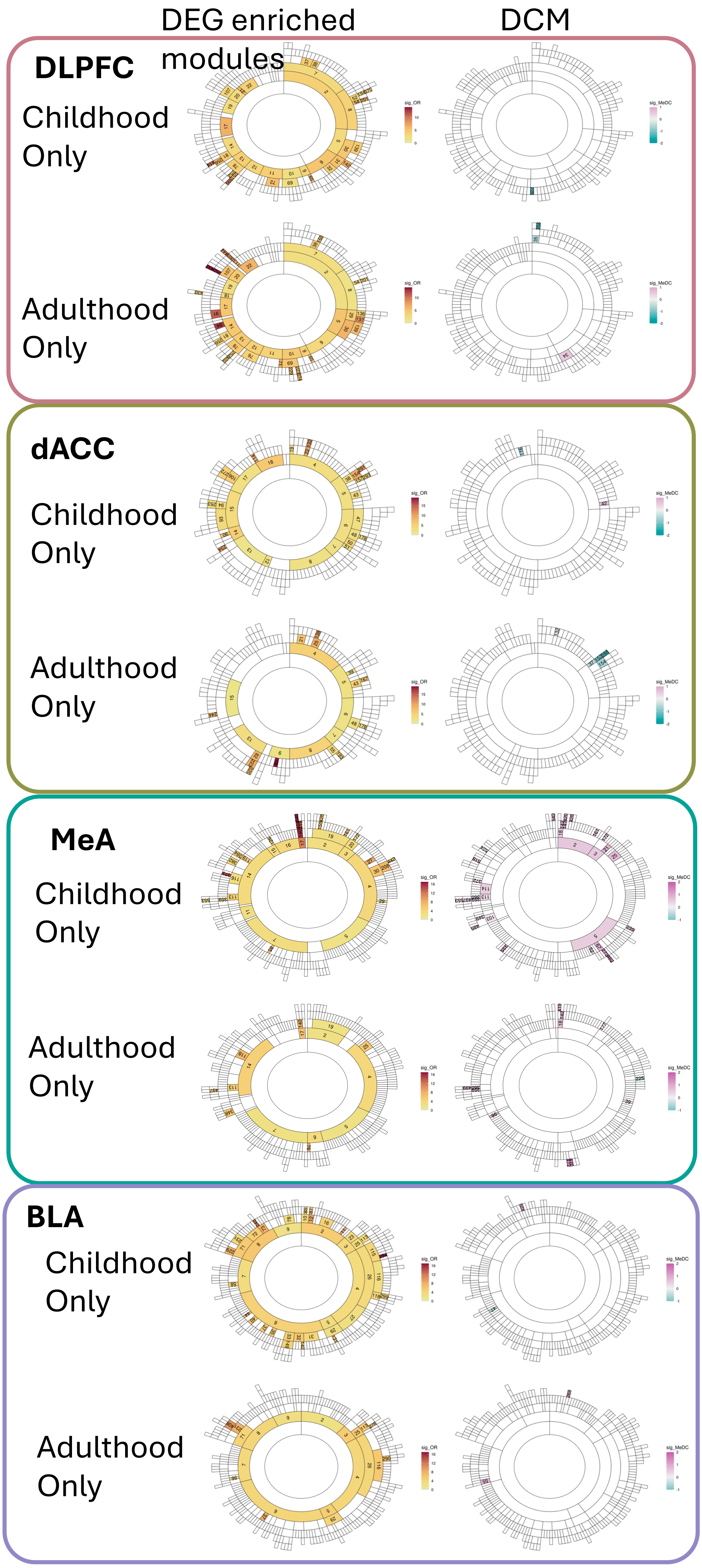

### SFig8

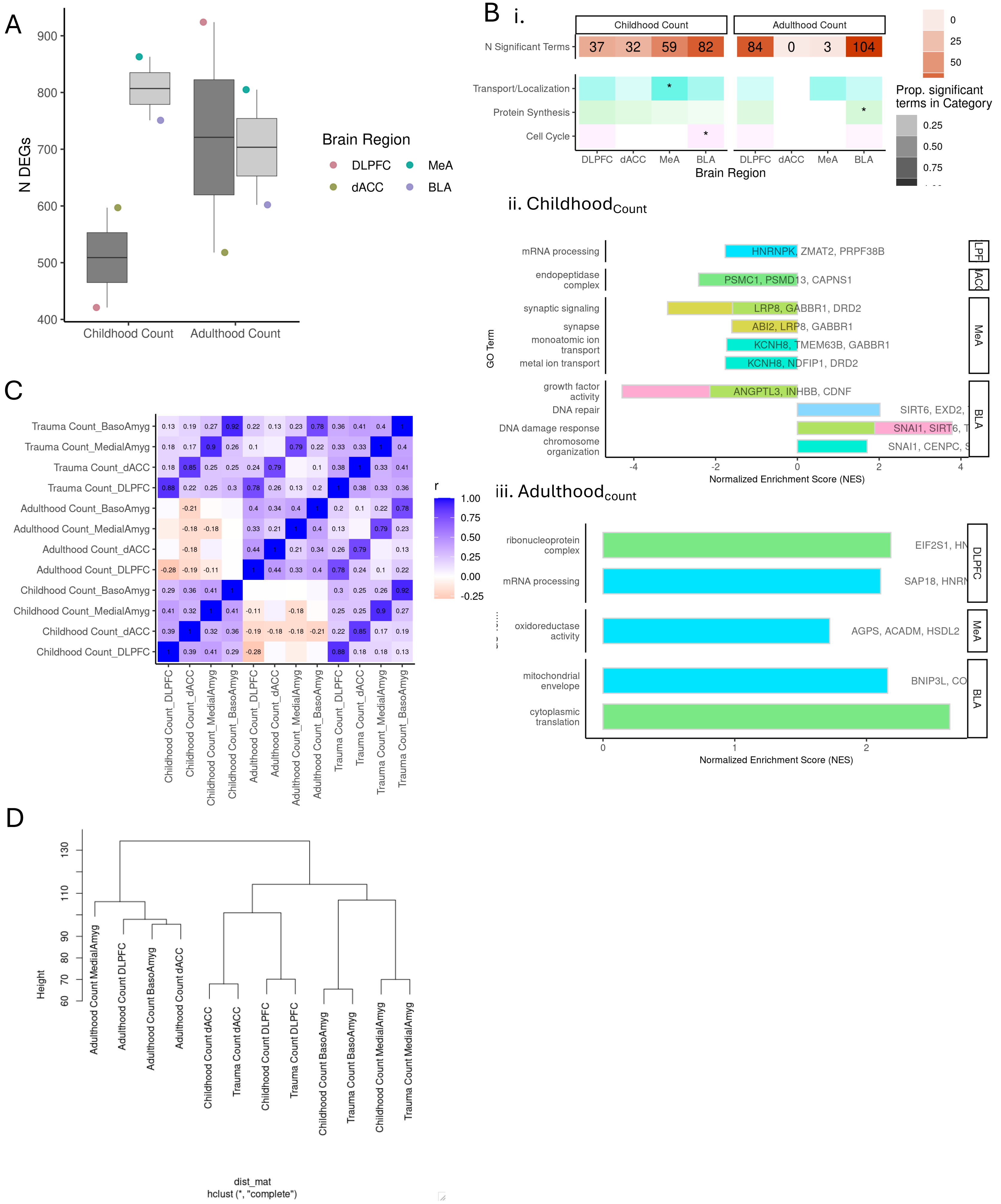

### SFig9

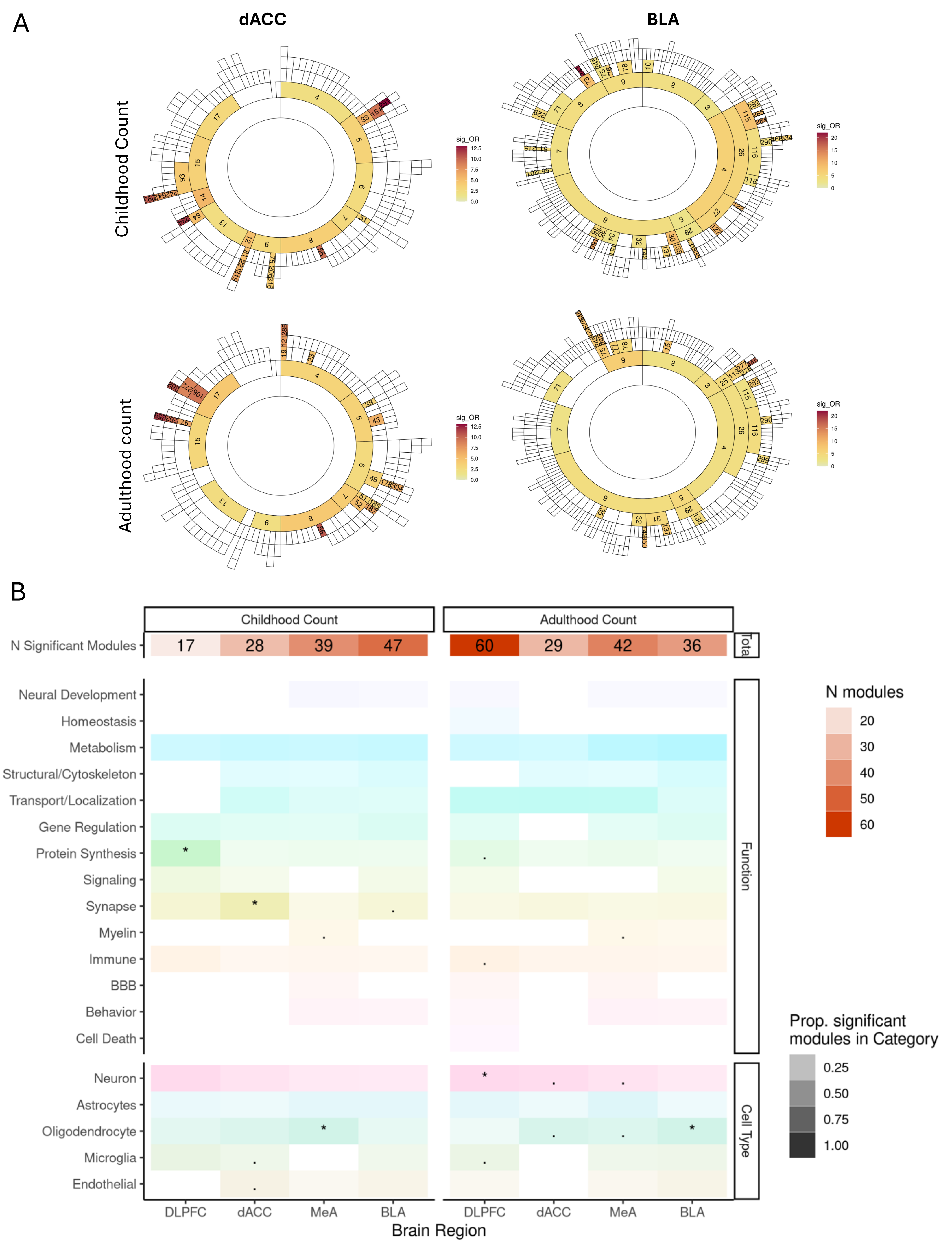

### SFig10

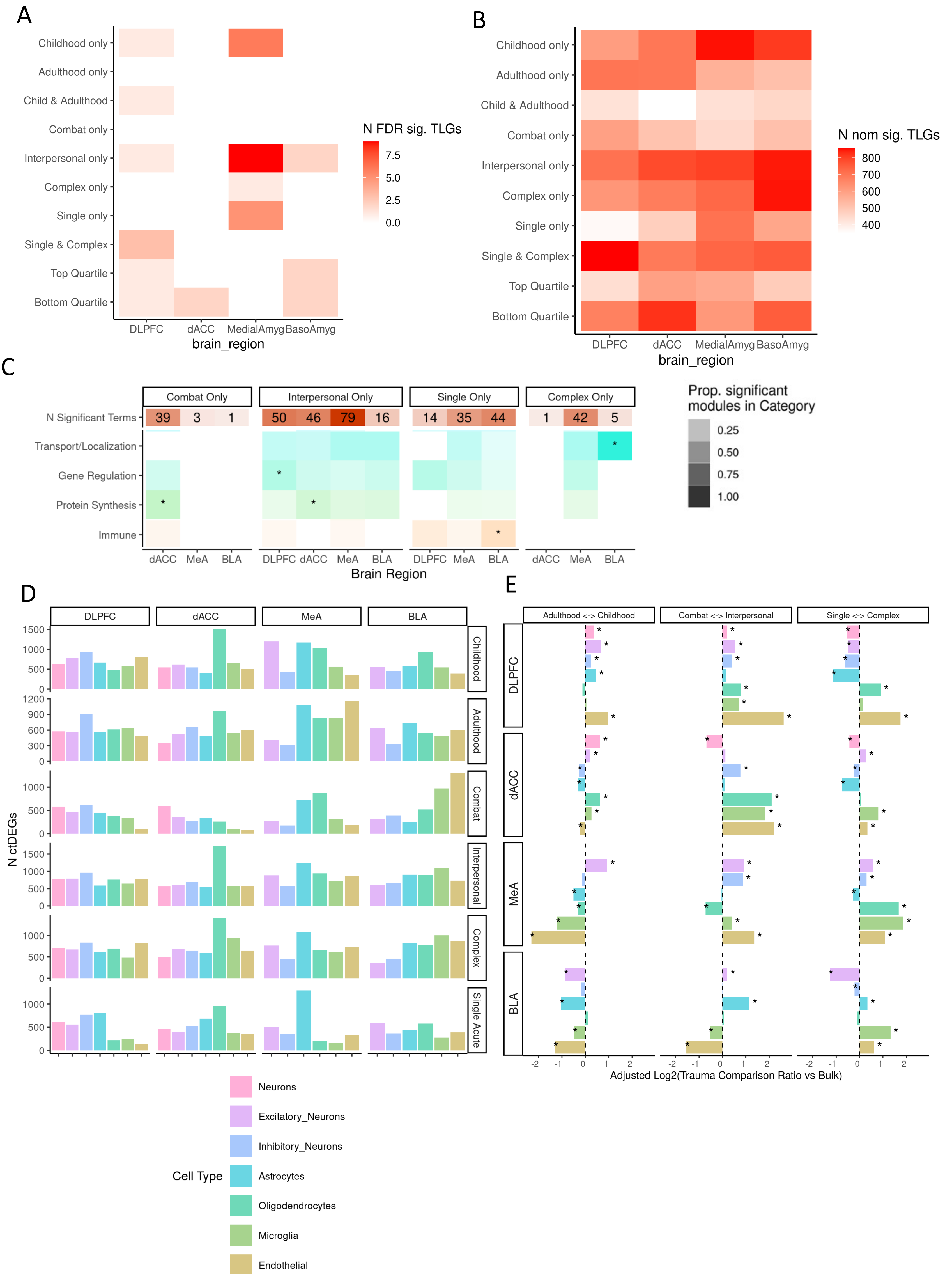

### SFig11

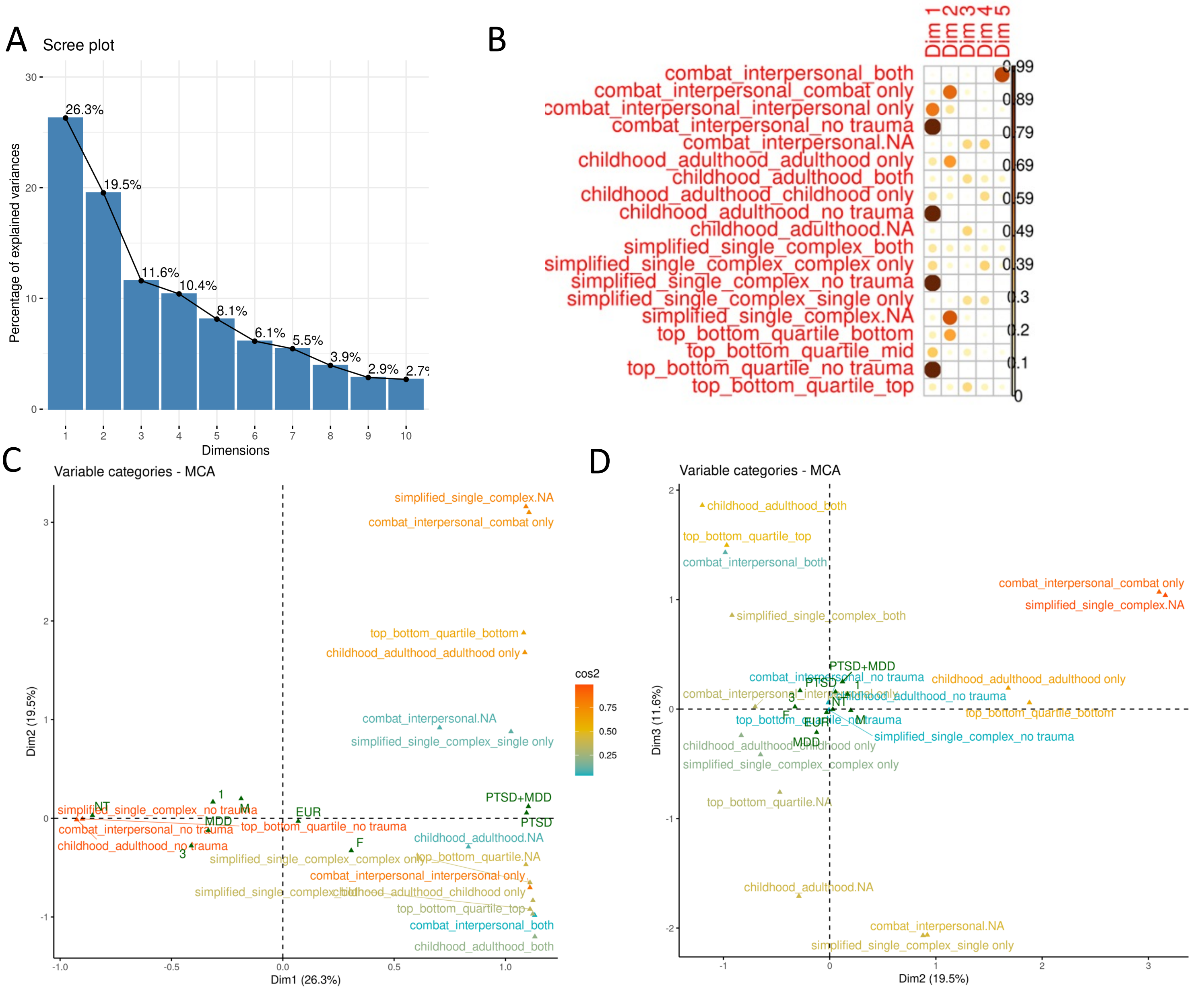

### SFig13

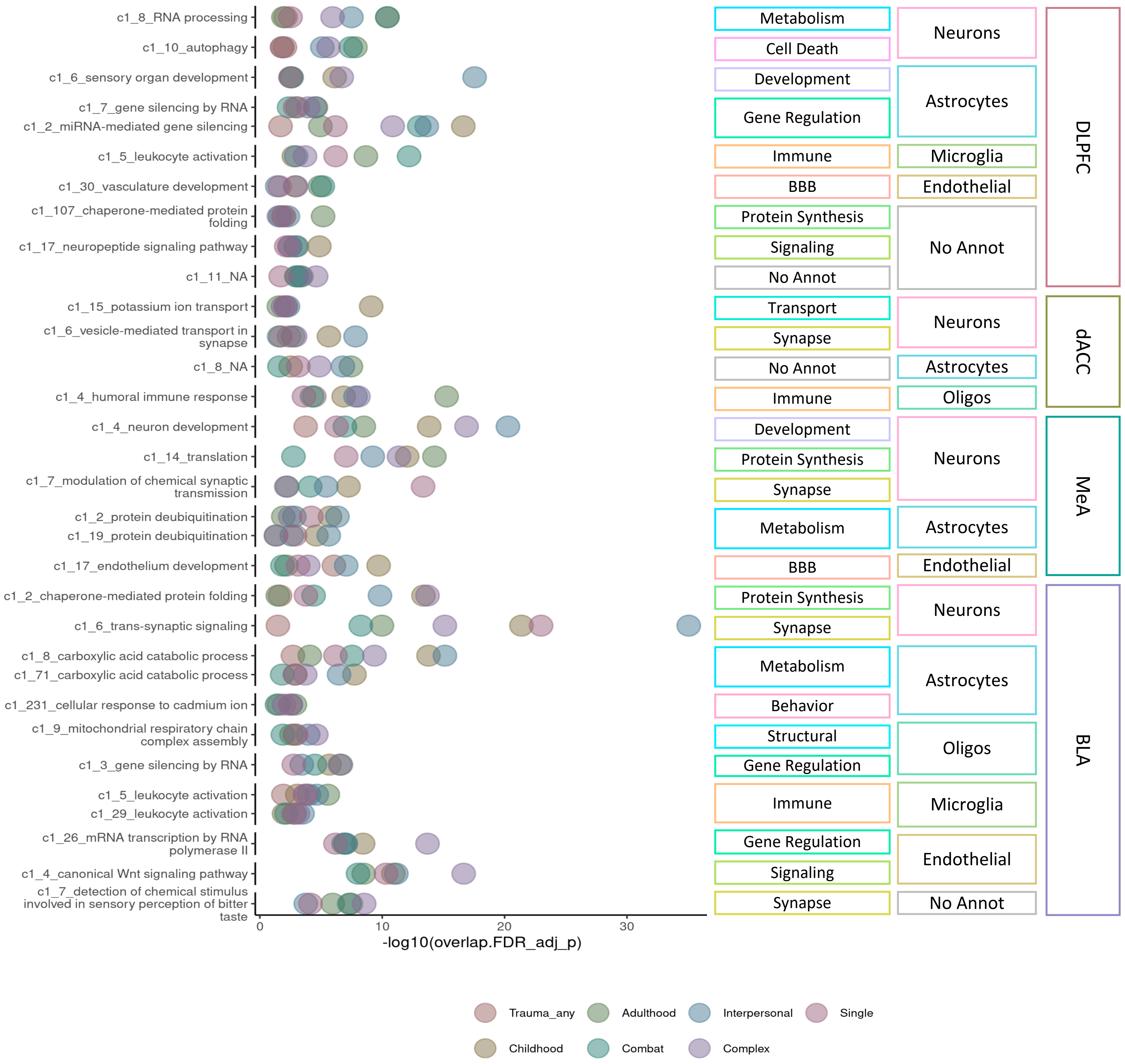

### SFig14

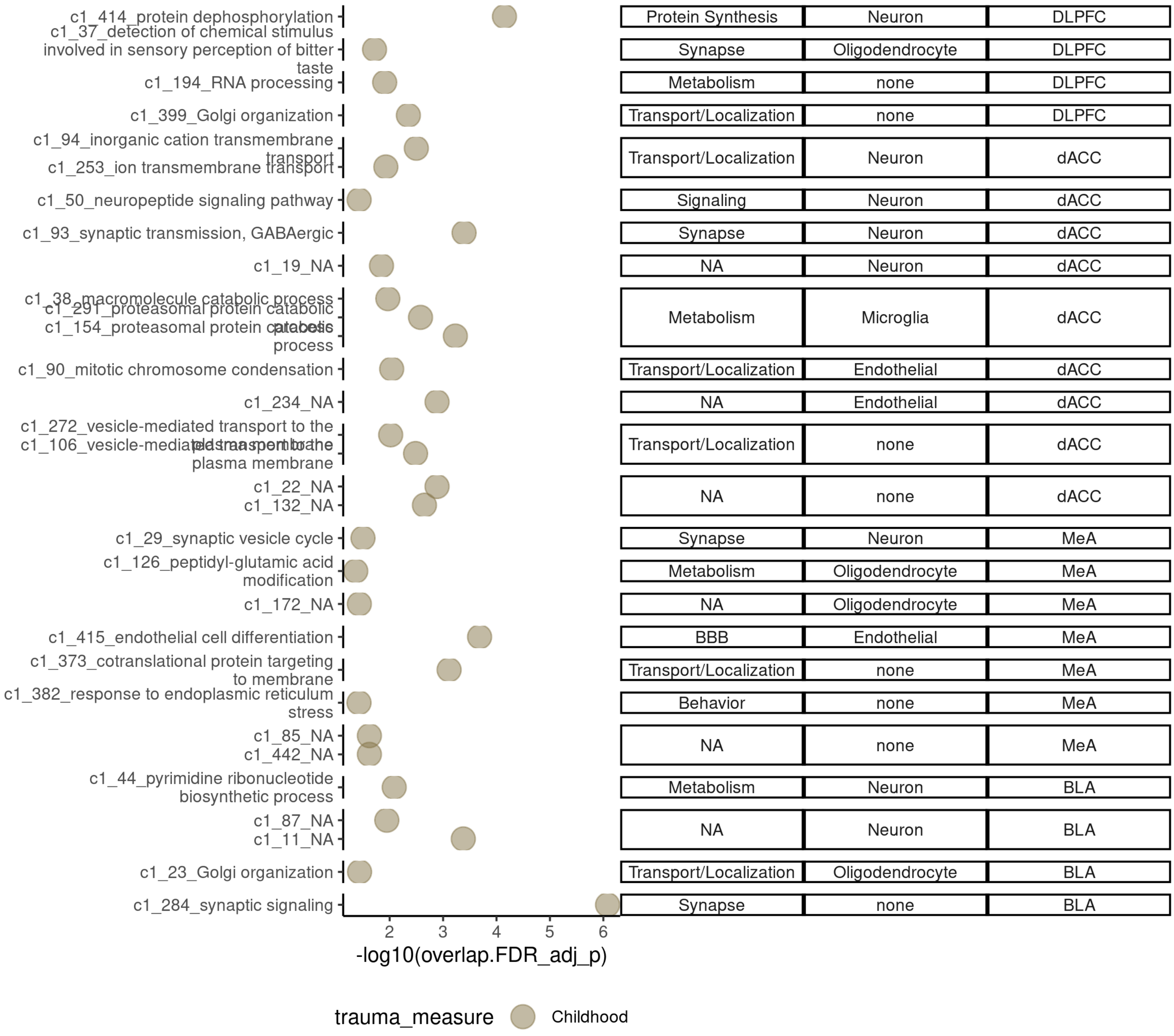

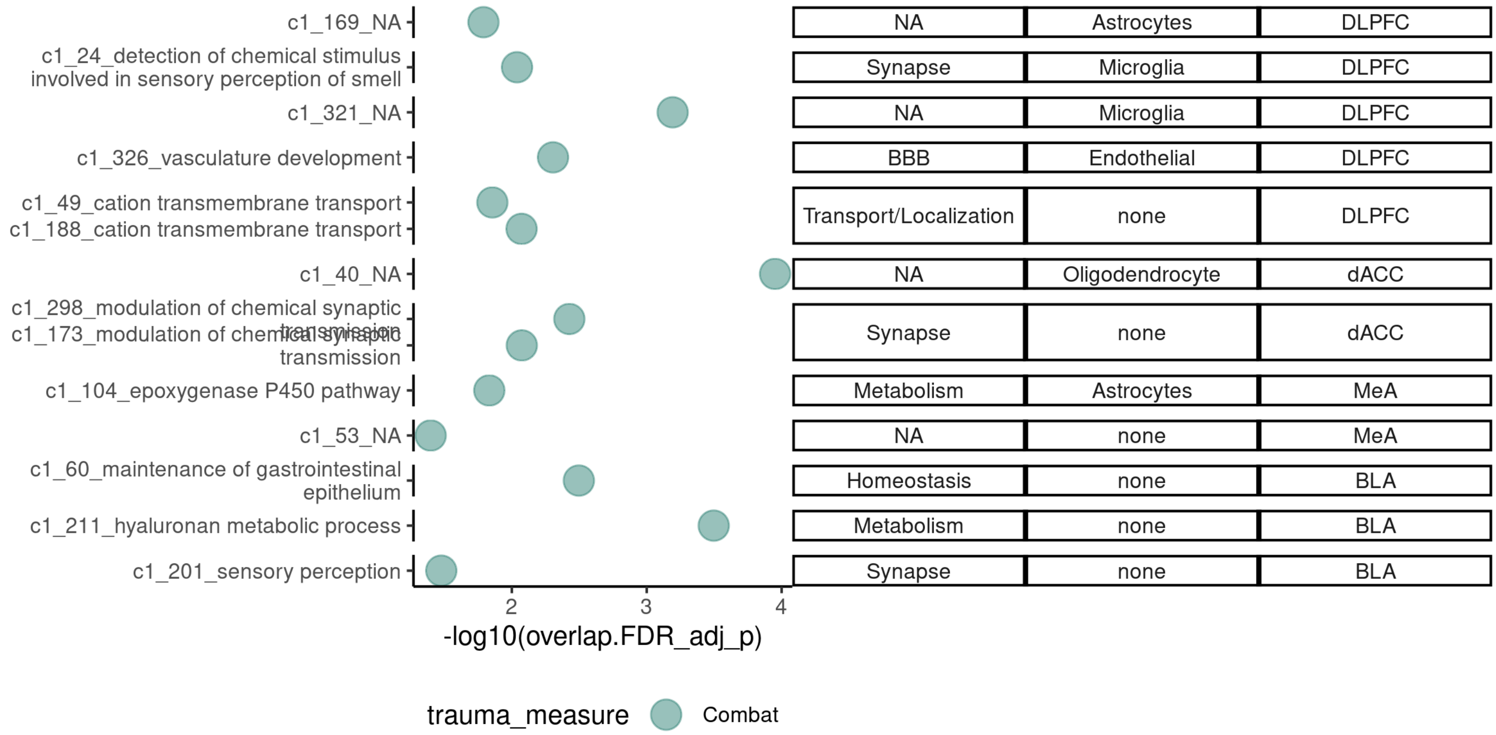

### SFig15

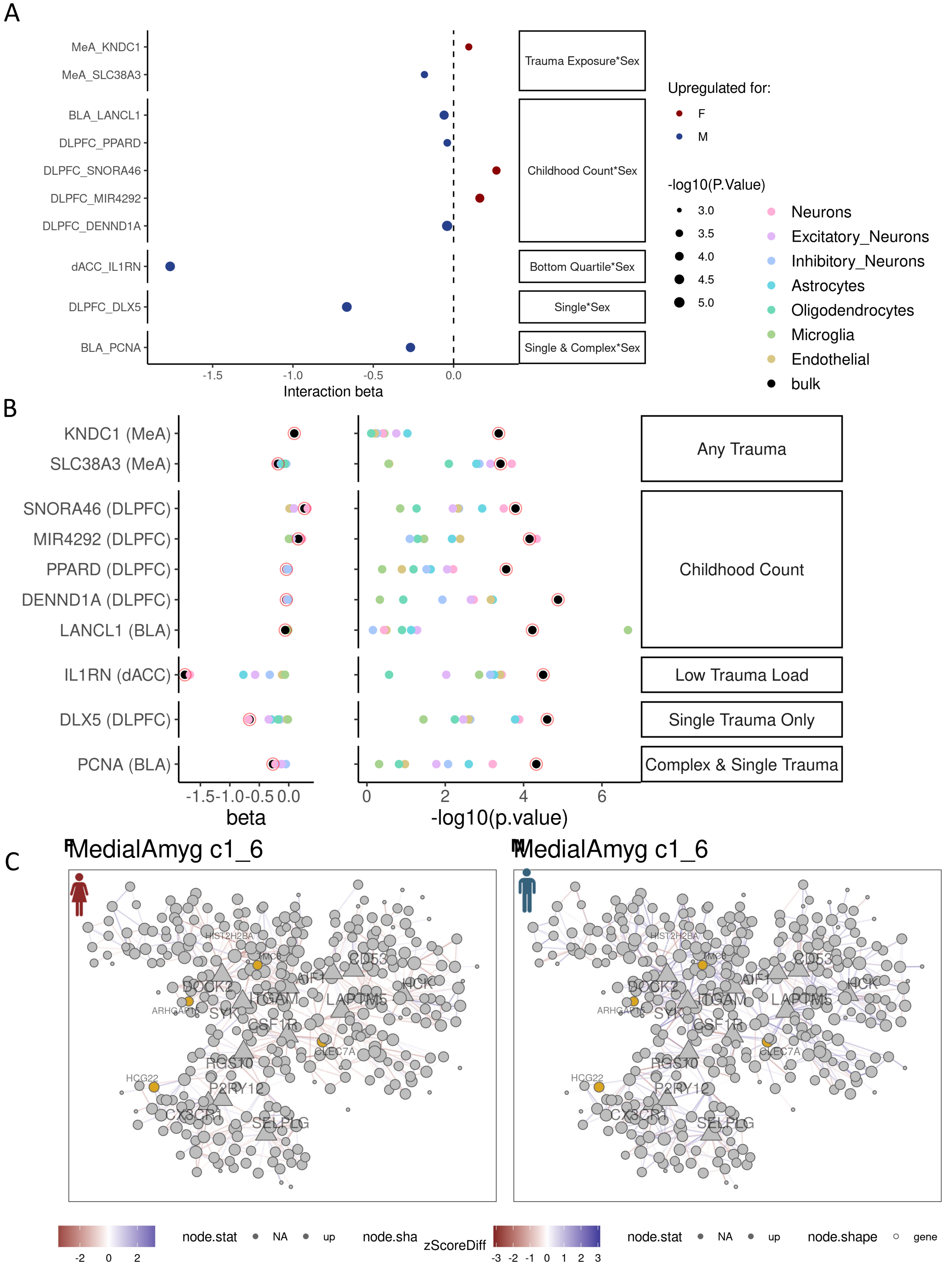

### SFig16

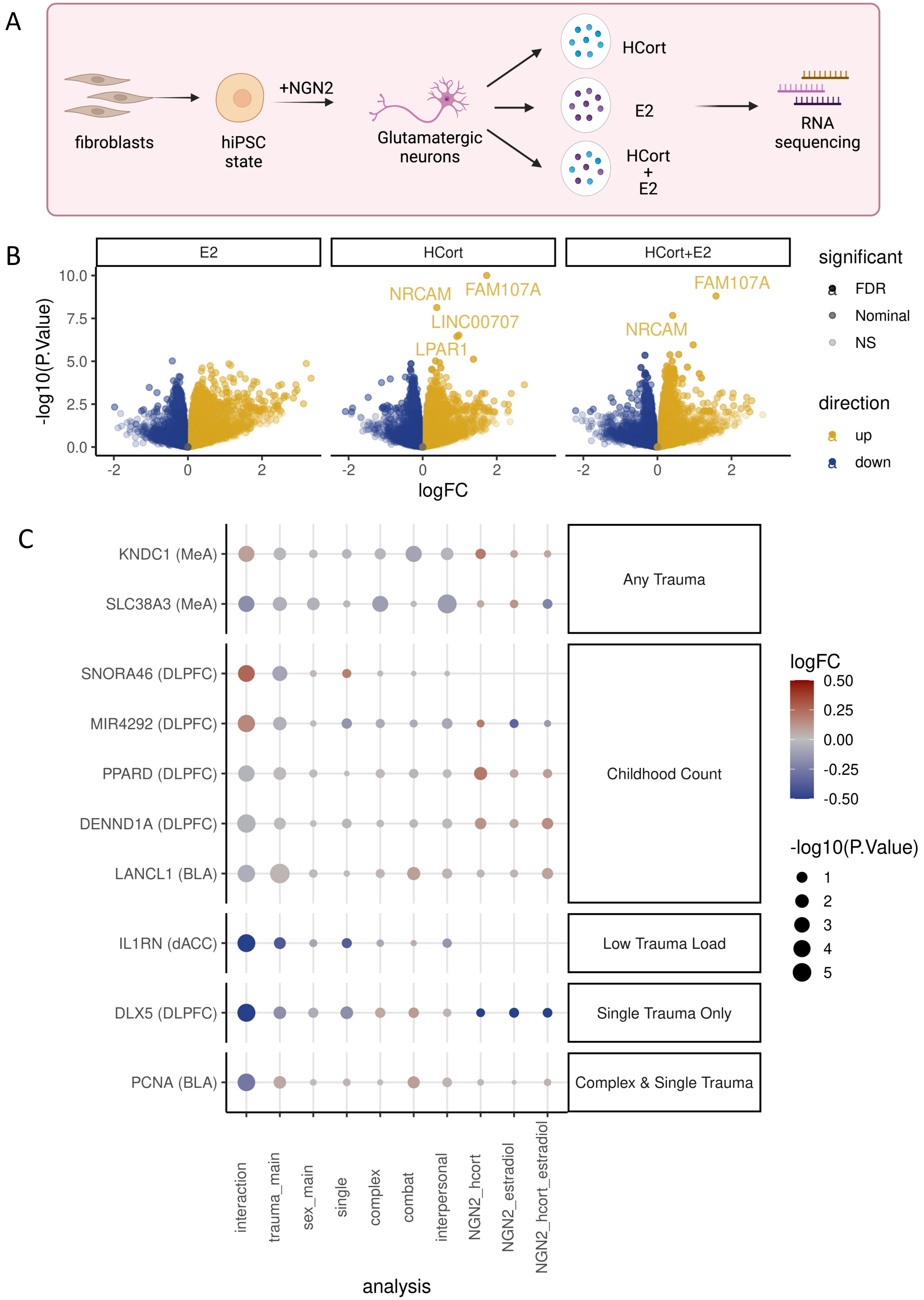
