## Supplementary material for "Decoding the transcriptomic signatures of psychological trauma in human cortex and amygdala": SFig12

Venn Diagram of Common Modules - DLPFC

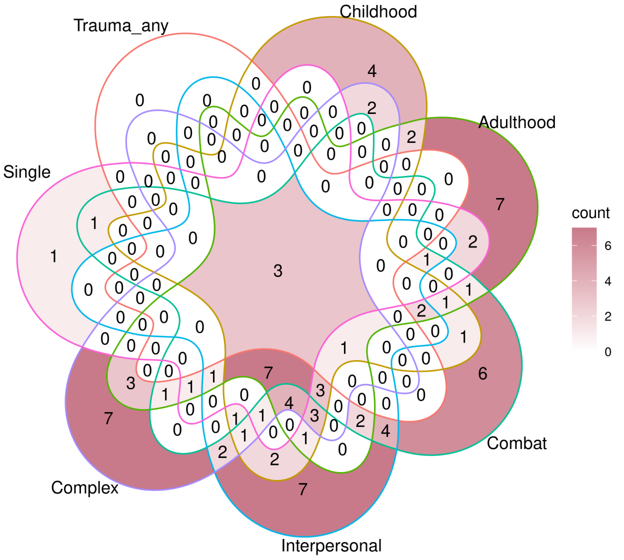


Venn Diagram of Common Modules - dACC

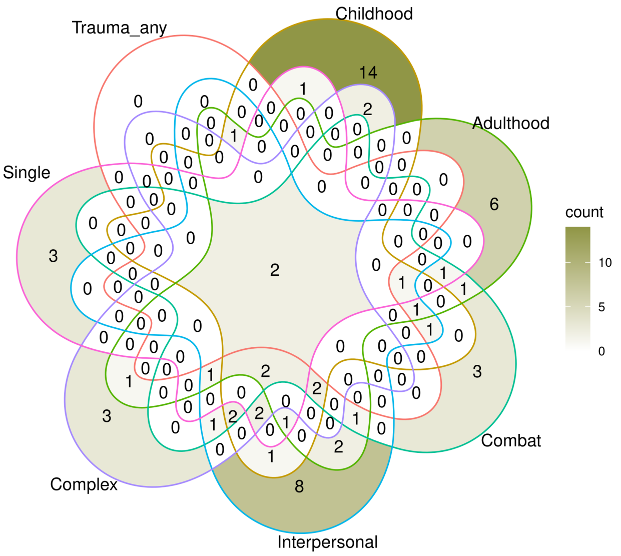


Venn Diagram of Common Modules - MedialAmyg

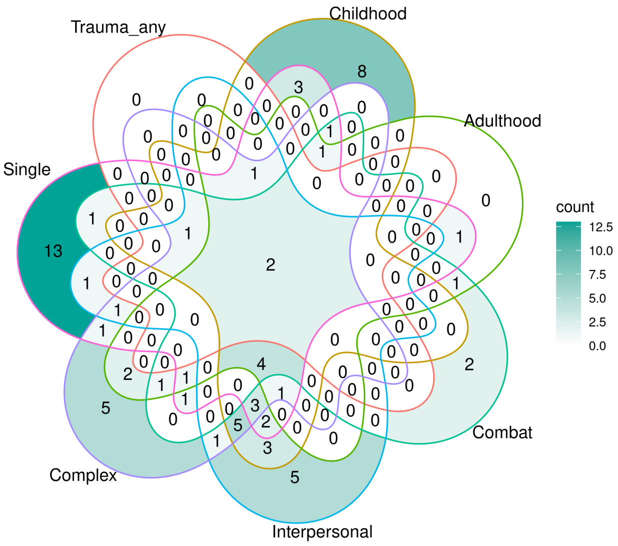


Venn Diagram of Common Modules - BasoAmyg

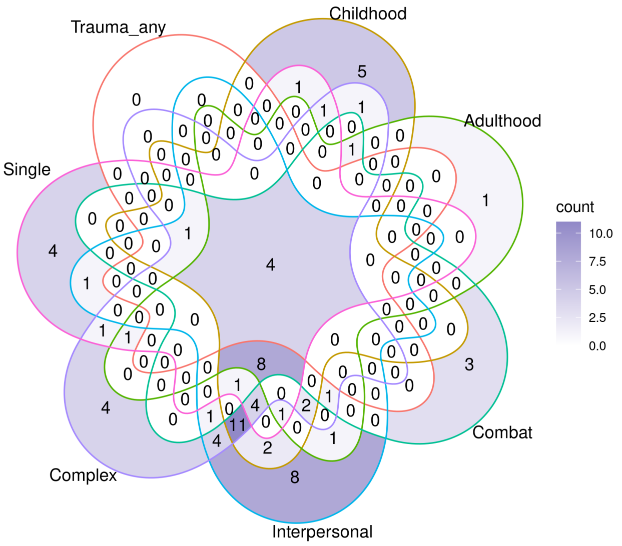
