## SupplementaaryText for "Decoding the transcriptomic signatures of psychological trauma in human cortex and amygdala"

**SUPPLEMENTAL TEXT**

Transcriptomic signatures of trauma are distinct from that of diagnosis

Trauma, diagnosis, and sex are highly interrelated, and individuals in our sample are enriched for diagnosis of MDD (35%) or PTSD (35%)(**SFig 1B**). We calculated PTSD and MDD DEGs, and tested for genome-wide correlation with trauma exposure and cumulative trauma DEGs (**SFig. 1C-D**). There is a relatively high correlation between transcriptional signatures of PTSD and trauma exposure (r=0.81-0.83) and to a lesser degree between PTSD and trauma_Count_ (r=0.64-0.68) across all four brain regions. We observed a small negative correlation between MDD and trauma exposure (r=-0.19--0.1) and trauma_Count_ (r=-0.12—0.245) .

Because of the degree of overlap in our sample, covarying out these factors is not possible. Instead, we performed a series of sensitivity analyses to determine whether significant associations were driven by trauma itself, or were instead associated more broadly with PTSD, MDD, or sex (**SFig 1A**). We tested all 6924 nominally significant trauma measure-brain region gene associations and found 3004 (43.3% of all tested associations) met criteria as a trauma-leading association (**SFig 1E-F**). In this study, we analyze only those gene associations for which trauma is a driving factor, termed trauma-leading genes.
